# Targeting Ribonucleotide Reductase Overcomes FLT3 Inhibitor Resistance in Acute Myeloid Leukemia

**DOI:** 10.64898/2026.07.21.736123

**Authors:** Zhen Tian, Hanghang Zhang, Srinivas Chatla, Xiaoyu Wei, Yang Liu, Dongwook Kim, Yahui Li, Peng Wang, Yun Liao, Xiaolei Liu, Dan Yang, Stacia Octaviani, Gui Ma, Anthony Pompetti, Gennaro Calendo, Martin P. Keough, Wei Xu, Jing Zhang, Hong Zheng, Elliot Stieglitz, Catherine Smith, Jian Huang

## Abstract

Internal tandem duplication mutations in *FLT3* (*FLT3*^ITD^) occur in approximately 30% of patients with acute myeloid leukemia (AML) and are among the most common genetic alterations in this disease. *FLT3*^ITD^ is a major driver of AML and is associated with poor clinical outcomes. Although FLT3 inhibitors (FLT3is) have significantly improved outcomes for patients with *FLT3*^ITD^+ AML, acquired resistance remains a major barrier to durable clinical benefit. Reactivation of RAS/MAPK signaling, often driven by activating *NRAS* mutations, is a major mechanism of FLT3i resistance in AML; however, effective strategies to overcome this resistance remain lacking. Here, we identify ribonucleotide reductase (RNR) as a critical therapeutic vulnerability in *NRAS*-driven FLT3i-resistant *FLT3*^ITD^+ AML. Activation of RAS signaling through *SPRY3* loss or oncogenic *NRAS* mutations confers robust resistance to FLT3is, whereas pharmacologic inhibition of RNR with multiple inhibitors, as well as siRNA-mediated RNR suppression, reverses FLT3i resistance and restores FLT3i sensitivity across multiple *FLT3*^ITD^+ AML models *in vitro*. *In vivo*, clofarabine, an FDA-approved RNR inhibitor (RNRi), significantly overcomes *NRAS* mutation-driven FLT3i resistance. In combination with FLT3 inhibition, clofarabine markedly suppresses the progression of FLT3i-resistant AML and significantly prolongs survival in cell line-derived xenograft (CDX) models. Importantly, the therapeutic efficacy of the gilteritinib/clofarabine combination was independently validated in two genetically distinct patient-derived xenograft (PDX) models harboring different *NRAS* mutations, demonstrating robust reduction of leukemia burden and confirming the generalizability of RNR inhibition in primary FLT3i-resistant AML. Together, these findings identify a previously unrecognized therapeutic vulnerability in FLT3i-resistant *FLT3*^mut^+ AML and establish RNR inhibition as an effective strategy to overcome FLT3i resistance, providing a strong rationale for the clinical evaluation of RNRis in combination with FLT3is in patients with resistant AML.

**Significance:** Although FLT3 inhibitors (FLT3i) are an important therapeutic advance in *FLT3*^ITD^+ AML, resistance commonly develops. We identified ribonucleotide reductase (RNR) as a new key vulnerability in *NRAS*-driven FLT3i-resistant AML and demonstrated that multiple RNRis, including the FDA-approved agent clofarabine, restore FLT3i sensitivity and enhance antileukemic activity, supporting a clinically actionable combination strategy.

## INTRODUCTION

Acute myeloid leukemia (AML) is an aggressive hematologic malignancy characterized by the uncontrolled proliferation and impaired differentiation of myeloid blasts, leading to bone marrow failure and high mortality. Despite advances in therapeutic strategies, outcomes remain poor, particularly in relapsed or refractory disease, where treatment resistance is common and 5-year survival rates are approximately 10%^1,2^.

Among the most frequently mutated genes in AML is Fms-like tyrosine kinase 3 (FLT3), with internal tandem duplication (ITD) mutations occurring in approximately 30% of patients^3,4^. *FLT3*^ITD^ drives constitutive activation of downstream signaling pathways, including RAS/MAPK/ERK and PI3K/Akt, promoting leukemic cell proliferation and survival^4,5^. The development of FLT3is, including first-generation agents such as midostaurin and lestaurtinib and second-generation inhibitors such as quizartinib (AC220) and gilteritinib, has significantly improved clinical outcomes in patients with *FLT3*-mutant AML. Several of these agents are now approved for clinical use, including gilteritinib and quizartinib, which received FDA approval in 2018 and 2023, respectively^6–9^. However, responses are often transient, and most patients eventually relapse due to acquired resistance, which significantly limits the efficacy of targeted AML therapies and remains a major clinical challenge^10,11^.

FLT3i resistance develops in approximately 50-60% of responding patients within 6-12 months of treatment and is driven by diverse molecular and cellular mechanisms^12,13^. A common mechanism arises from secondary mutations in the FLT3 tyrosine kinase domain (TKD), including F621L, A627P, F691L, and Y842C, which alter the activation loop and lead to constitutive kinase activation and resistance to inhibitors targeting *FLT3*^ITD^ mutations^10,11^. Beyond on-target mutations, AML cells can acquire resistance through reactivation of pro-survival signaling pathways, particularly the RAS/MAPK cascade, which is observed in approximately one-third of patients with *FLT3*-mutated AML who develop FLT3i resistance^14,15^. RAS proteins are small GTPases that function as molecular switches regulating cell proliferation, survival, and growth, and mutations in RAS family members, including *NRAS* and *KRAS*, are among the most common oncogenic events in AML^16^. Notably, unlike in solid tumors where *RAS* mutations often occur early, *NRAS* mutations in AML are frequently acquired during disease progression or following therapy, particularly in relapsed or refractory settings^17,18^. *NRAS* mutations, present in approximately 10–15% of AML cases and commonly affecting codons 12, 13, and 61, result in constitutive activation of downstream MAPK signaling^19^. In the context of *FLT3*-mutant AML, oncogenic *NRAS* mutations (e.g., *NRAS*^G12C^ and *NRAS*^Q61K^) have been strongly associated with resistance to FLT3is, enabling leukemic cells to sustain proliferative and survival signaling despite FLT3 blockade^18,20^. Consistent with this, our previous CRISPR-based genetic screen identified *SPRY3* as a novel regulator of FLT3i sensitivity. We found that loss of *SPRY3*, a negative regulator of RAS/MAPK signaling, activates downstream MAPK signaling and confers robust resistance to FLT3is in AML, suggesting that *SPRY3* loss and oncogenic *NRAS* mutations converge on a shared RAS/MAPK signaling axis to drive FLT3i resistance^21^. However, the mechanisms by which RAS/MAPK pathway activation is translated into downstream programs that sustain leukemic proliferation under FLT3 inhibition and promote FLT3i resistance remain poorly defined.

Ribonucleotide reductase (RNR), the rate-limiting enzyme in deoxyribonucleotide (dNTP) synthesis, is essential for DNA replication and repair by controlling the availability of dNTP pools required for genome maintenance^22,23^. RNR functions as a heteromeric complex composed of RRM1 and RRM2 catalytic subunits, as well as the stress-inducible subunit RRM2B, enabling cells to adapt nucleotide production under conditions of replication stress or DNA damage^23,24^. Consistent with its central role in supporting DNA synthesis and cell proliferation, elevated RNR activity has been implicated in therapeutic resistance across multiple cancer types^25,26^. Notably, several established RNRis, including clofarabine, fludarabine, gemcitabine, and hydroxyurea are already FDA-approved drugs^27^. However, RNR has not previously been recognized as a critical downstream mediator of NRAS-driven FLT3i resistance in *FLT3*^ITD^+ AML.

Here, we identify RNR as a key downstream effector of RAS signaling in FLT3i resistance. We show that loss of *SPRY3* or *NRAS* mutations drives FLT3i resistance through activation of an RNR-dependent nucleotide metabolic program in AML. Targeting RNR restores sensitivity to FLT3i *in vitro* and *in vivo*, including in both cell line-derived xenograft (CDX) and two genetically distinct patient-derived xenograft (PDX) models of AML. Together, these findings uncover a previously unrecognized link between RAS signaling and nucleotide metabolism in AML and support RNR inhibition, including repurposing clinically available agents such as clofarabine, as an effective therapeutic strategy to overcome FLT3i resistance in *FLT3*^ITD^+ AML.

## METHODS

### Cell lines

MV4-11 *SPRY*WT and *SPRY*KO cell lines were generated as previously described^21^. Doxycycline-inducible MV4-11 and MOLM-14 cell lines harboring *NRAS*^WT^, *NRAS*^G12C^, and *NRAS*^Q61K^ mutations were kindly provided by Dr. Catherine C. Smith (University of California, San Francisco, CA) and established as previously described^15^. MOLM-14 cells expressing mCherry, including *NRAS*^WT^ and *NRAS*^G12C^ variants, were kindly provided by Dr. Elliot Stieglitz (University of California, San Francisco, CA). MV4-11 cells were cultured in IMDM supplemented with 10% FBS and 1% penicillin/streptomycin/L-glutamine. MOLM-14 cells were maintained in RPMI-1640 medium supplemented with 10% FBS and 1% penicillin/streptomycin/L-glutamine. All cell lines were routinely tested and confirmed to be free of mycoplasma contamination.

### Patient samples

Patient samples for PDX model #1 were kindly provided by Elliot Stieglitz (University of California, San Francisco, CA). Primary AML specimens were obtained through institutional tissue banking protocols at the University of California, San Francisco under Institutional Review Board-approved protocols. All samples were de-identified prior to use. Written informed consent was obtained from all patients for the collection and research use of their specimens in accordance with the Declaration of Helsinki. Paired samples from the same patient, collected at diagnosis and at relapse following gilteritinib treatment^38^, were used in this study. Patient samples for PDX model #2 were kindly provided by the Stem Cell and Xenograft Core, Perelman School of Medicine, University of Pennsylvania (RRID: SCR_010035). All human specimens were obtained under Institutional Review Board-approved protocols with written informed consent from patients in accordance with the Declaration of Helsinki.

### Compounds

Gilteritinib (HY-12432), AC220 (quizartinib, HY-13001), trametinib (HY-10999), clofarabine (HY-A0005), fludarabine (HY-B0069), gemcitabine (HY-17026), hydroxyurea (HY-B0313), cladribine (HY-13599), triapine (HY-10082), nelarabine (HY-13701), doxycycline (HY-N0565), and D-luciferin potassium (HY-12591B) were purchased from MedChemExpress (MCE).

### Cell viability assays

Cells were seeded at a density of 1 × 10^4^ cells per well in three technical replicate 96-well plates and treated as indicated. For doxycycline-inducible MV4-11 and MOLM-14 cell lines expressing mutant *NRAS*, cells were pretreated with doxycycline (1 μg/mL) for 24 hours before the experiments and maintained in doxycycline-containing medium throughout the duration of the experiments. Cell viability was assessed using the Cell Counting Kit-8 (CCK-8; HY-K0301, MCE) according to the manufacturer’s instructions.

### RAS Activation Assay

RAS activity was measured using the Ras GTPase Chemi ELISA Kit (Active Motif, Cat. No. 52097) according to the manufacturer’s instructions. Briefly, cells were lysed and whole-cell lysates were collected. Protein concentrations were determined using a BCA assay (Thermo Scientific, #A55864). Equal amounts of protein lysates were incubated in glutathione-coated 96-well plates preloaded with GST-Raf-RBD, which selectively binds GTP-bound (active) RAS. Following washing, bound RAS was detected using an anti-RAS primary antibody and an HRP-conjugated secondary antibody. Chemiluminescent substrate was then added, and luminescence was measured using a microplate luminometer. RAS activity was expressed as relative luminescence units (RLU).

### Real-time PCR

Total RNA Total RNA was extracted from harvested cells using TRIzol reagent (Invitrogen, #15596026). cDNA was synthesized from 1 µg of total RNA using High-Capacity cDNA Reverse Transcription Kit (Applied Biosystems, #3154782). Quantitative PCR was performed using Power SYBR Green Master Mix (Applied Biosystems, 4367659) on a QuantStudio 6 Flex Real-Time PCR System (Applied Biosystems). Sequence primers for target genes, including *RRM1*, *RRM2*, *RRM2B*, are described in the “List of primers” section. Thermal cycling conditions were as follows: 95°C for 2 minutes, followed by 40 cycles of 95°C for 15 seconds and 60°C for 1 minute. Relative gene expression was calculated using the 2^^-ΔΔCt^ method, with GAPDH as the endogenous control.

### Small interfering RNAs (siRNAs)-mediated gene knockdown

Transient gene knockdown was performed using siRNAs and TransIT-TKO® Transfection Reagent (Mirus Bio, #MIR 2150) according to the manufacturer’s instructions. Briefly, cells were transfected with either control or gene-specific siRNAs including siRRM1 (MCE, # HY-RS12285), siRRM2 (MCE, #HY-RS12286), siRRM2B (MCE, #HY-RS12287), siE2F1 (MCE, #HY-RS04110), or siMYC (MCE, #HY-RS08877), and cultured under standard growth conditions. Twenty-four hours after siRNA transfection, cells were treated with the indicated drugs while maintaining siRNA transfection conditions. Cells were subsequently analyzed for downstream assays, and knockdown efficiency was confirmed by RT-qPCR and immunoblotting.

### Western blot

Cells were plated in appropriate media under the indicated treatments. After 48 h incubation, cells were washed with PBS and lysed in RIPA lysis buffer supplemented with protease and phosphatase inhibitors. Lysates were clarified by centrifugation, and protein concentrations were determined using a BCA assay (Thermo Scientific, #A55864). Equal amounts of protein were separated by Bis-Tris gel electrophoresis and transferred onto nitrocellulose membranes using the iBlot 3 system (Invitrogen). Membranes were blocked with 5% nonfat milk and incubated overnight at 4°C with primary antibodies including anti-RRM1 (Proteintech, #10526-1-AP), anti-RRM2 (Proteintech, #11661-1-AP), anti-RRM2B (Proteintech, #18005-1-AP), anti-E2F1 (Cell signaling Technology, #3742S), and anti-c-Myc (Cell signaling Technology, #5605T). After incubation with HRP-conjugated Protein A secondary antibody (MilliporeSigma, #18-160), protein bands were visualized using enhanced chemiluminescence (ECL) on the iBright imaging system (Invitrogen).

### Cell proliferation and apoptosis assays

Cells were seeded in 12-well plates at a density of 2 × 10⁵ cells per well and treated with vehicle, gilteritinib, clofarabine, or their combination for 48 h. After treatment, cells were harvested and washed twice with cold PBS, then fixed and permeabilized. For proliferation analysis, followed by intracellular staining with Brilliant Violet 421 anti-mouse Ki-67 antibody (BioLegend, #350505). For apoptosis analysis, cells were stained with FITC Annexin V (Biolegend, #640906). Flow cytometry data were analyzed with FlowJo.

### Cell cycle analysis

AML cells were harvested 24 h after drug treatment, fixed overnight in pre-chilled 70% ethanol at 4°C, and stained with propidium iodide (PI) buffer containing RNase A. DNA content was analyzed using a Sony flow cytometer, and the percentages of cells in G0/G1, S, and G2/M phases were quantified using the manufacturer’s analysis software.

### Intracellular dNTP quantification

Intracellular deoxyribonucleotide triphosphate (dNTP) levels were quantified using a PCR-based primer extension assay as previously described^37^. Briefly, following the indicated treatments, 5 × 10^6^ cells were harvested and resuspended in ice-cold 60% methanol. Cell extracts were vortexed vigorously, boiled for 3 min, and centrifuged at 16,000 × g for 20 min at 4°C. Supernatants were collected and filtered through pre-equilibrated Amicon Ultra 0.5-mL centrifugal filters (Millipore). Filtrates were dried using a vacuum centrifuge at 70°C, and the resulting pellets were resuspended in nuclease-free water and stored at −80°C until analysis. dNTP levels were measured using a DNA polymerase-based primer extension assay performed on Real-Time PCR System. Reaction mixtures contained primer, probe, and template at final concentrations of 0.4 μM each, MgCl₂ at 2 mM, and excess non-limiting dNTPs at 100 μM, excluding the dNTP being measured. AmpliTaq Gold DNA polymerase was added at 0.875 U per reaction in a final reaction volume of 25 μL containing 2.5 μL of cell extract. Reactions were initiated with a 10-min incubation at 95°C followed by primer extension at 60°C for up to 30 min. Fluorescence signals were collected throughout the extension reaction and analyzed after subtraction of blank controls lacking the limiting dNTP. Intracellular dNTP levels were normalized to GTP levels and expressed as dATP/GTP, dGTP/GTP, and dCTP/GTP ratios.

### BrdU incorporation assay

DNA synthesis and proliferation were assessed using the Fluorescent BrdU Assay Kit (BiCell Scientific, #BCBRDU). Cells were treated with vehicle, gilteritinib, clofarabine, or their combination for 48 h, followed by BrdU labeling (1:1000 dilution) for 30 min at 37°C with 5% CO_2_. Cells were washed with PBS, fixed with cold methanol for 30 min at −20°C, air-dried, and blocked with antibody block solution. Cells were then incubated with anti-BrdU primary antibody (1:50) for 2 h, followed by FITC-conjugated secondary antibody (1:200) for 1 h at room temperature. After washing, slides were mounted with Mowiol solution and imaged by fluorescence microscopy. BrdU-positive cells were quantified and normalized to the vehicle-treated group.

### Cell-derived xenograft models

All *in vivo* xenograft studies were performed in accordance with protocols approved by the Institutional Animal Care and Use Committee (IACUC) of Cooper University Hospital. AML cells were transduced with a firefly luciferase-expressing lentiviral construct and injected into NSG (Nod.Cg-Prkdc^scid^Il2rg^tm1Wjl^/SzJ) immunocompromised mice. For MV4-11 *SPRY3*WT or *SPRY3*KO CDX models, 1 × 10^6^ cells were injected, for *NRAS*^WT^ and *NRAS*^G12C^ CDX models, 2.5 × 10^6^ cells were injected. After engraftment of AML was confirmed by measuring the bioluminescence (BLI) using Ami imaging system (Spectral Instruments Imaging), mice were randomized into four treatment groups: vehicle, gilteritinib (30 mg/kg, po, qod), clofarabine (40 mg/kg, ip, qod), or the combination. Treatments were administered three times per week for a total of four doses. After completed administration, and leukemic burden was monitored by BLI using total flux in the unit of photons/second. Body weights were recorded throughout the study, and the body weight at the experimental endpoint was expressed as a percentage of the initial body weight. Mice were monitored daily for signs of disease progression, and overall survival was recorded. Survival curves were generated using the Kaplan–Meier method.

### Patient-derived xenograft models

For PDX model #1, patient-derived xenograft (PDX) studies were performed using female NSG mice in accordance with protocols approved by the Institutional Animal Care and Use Committee (IACUC) of Cooper University Hospital. Primary *FLT3*^ITD^-positive AML samples were obtained from a patient enrolled in a clinical trial of single-agent gilteritinib for relapsed or refractory FLT3-mutated AML. Paired samples were collected at diagnosis (gilteritinib-sensitive) and after relapse following gilteritinib treatment (gilteritinib-resistant). AML cells were transplanted into NSG mice via tail vein injection at a dose of 1 × 10⁶ cells per mouse. Leukemia engraftment was monitored by flow-cytometric analysis of human CD45 and CD33 expression in peripheral blood, and mice were considered engrafted when hCD45^+^/hCD33^+^ cells exceeded 1% of peripheral blood leukocytes. For leukemia burden studies, engrafted mice were randomized into four treatment groups (n = 5 per group) and treated with vehicle, gilteritinib (30 mg/kg, oral gavage, every other day), clofarabine (40 mg/kg, intraperitoneal injection, every other day), or the combination. Mice received a total of six doses of treatment. At study endpoint or upon reaching predefined humane endpoints, mice were euthanized and leukemia burden in bone marrow, spleen, and peripheral blood was quantified by flow-cytometric analysis of hCD45^+^/hCD33^+^ cells. Spleens were collected for weight measurement and histological analyses. Body weight at the study endpoint was recorded and expressed as a percentage of the initial body weight. For survival studies using the gilteritinib-resistant PDX model, engrafted mice were randomized into treatment groups (n = 8 per group). Mice receiving gilteritinib or clofarabine monotherapy continued treatment until reaching predefined humane endpoints. Mice receiving combination therapy were treated with gilteritinib and clofarabine for 12 weeks, after which treatment was discontinued and animals were monitored for survival. Mice were monitored for signs of disease progression, and overall survival was analyzed using the Kaplan–Meier method.

For the PDX model#2, Patient-derived xenograft (PDX) studies were performed using female NSG mice in accordance with protocols approved by the Institutional Animal Care and Use Committee at the University of Pennsylvania. Mice (6–9 weeks old) were conditioned with busulfan (25 mg/kg) 24 hours before intravenous injection of 5 × 10⁶ primary AML cells. Leukemia engraftment was confirmed by flow cytometric analysis of bone marrow aspirates 6 weeks after transplantation. Mice were then randomized into four treatment groups with comparable engraftment levels. Starting approximately 8 weeks after transplantation, mice received vehicle, clofarabine (40 mg/kg, intraperitoneally), gilteritinib (30 mg/kg, oral gavage), or the combination every other day for 2 weeks. At the end of treatment, bone marrow, spleen, and peripheral blood were collected, and leukemic burden was assessed by flow cytometry.

### Immunohistochemical (IHC) staining

Spleen tissues were fixed in 10% neutral-buffered formalin, paraffin-embedded, and sectioned. Tissue sections were deparaffinized, rehydrated, and subjected to antigen retrieval using Tris-based Antigen Unmasking Solution (Vector Laboratories, Burlingame, CA, USA). Following blocking with 10% normal serum, sections were incubated overnight at 4°C with an anti-human CD45 (hCD45) antibody (1:100; Thermo Fisher Scientific, #14945782). After washing with PBS, sections were incubated with biotinylated secondary antibodies for 30 min at room temperature. Antibody binding was detected using a streptavidin–peroxidase detection system with 3,3′-diaminobenzidine (DAB) as the chromogen (Vectastain ABC Kit, Vector Laboratories). Sections were counterstained with hematoxylin to visualize nuclei, dehydrated, mounted, and imaged using a bright-field microscope.

### Bulk RNA-seq

mRNA isolation, quality check, library construction, and sequencing were performed according to same protocol used previously^49^. All RNA-seq sequencing data are available at GEO (series accession number GSE339322).

### List of primes

RRM1: 5’-AAAGGAAGAGCAGCGTGCCAGA-3’ F; 5’-ACCTCATCCAGACCAGGACACT-3’ R RRM2: 5’-CTGGCTCAAGAAACGAGGACTG-3’ F; 5’-CTCTCCTCCGATGGTTTGTGTAC-3’ R RRM2B: 5’-ACTTCATCTCTCACATCTTAGCCT-3’ F; 5’-AAACAGCGAGCCTCTGGAACCT-3’ R GAPDH: 5’-GTCTCCTCTGACTTCAACAGCG-3’ F; 5’-ACCACCCTGTTGCTGTAGCCAA-3’ R

## RESULTS

### *SPRY* deficiency drives FLT3i resistance and activates RAS signaling in *FLT3*^ITD^+ AML

Previously, we identified loss of *SPRY3* as a top hit conferring resistance to the FLT3i AC220 in an unbiased genome-wide CRISPR screen^21^. SPRY3 belongs to the Sprouty family, which includes four members: SPRY1, SPRY2, SPRY3, and SPRY4^28^. We further found that individual knockout of *SPRY1*, *SPRY2*, or *SPRY3* conferred marked resistance to both AC220^21^ and gilteritinib (Fig. 1A), two FDA-approved FLT3 inhibitors. Because Sprouty proteins are established antagonists of receptor tyrosine kinase (RTK)-mediated RAS signaling^29^, we reasoned that *SPRY* loss may confer FLT3i resistance through activation of the RAS pathway. Supporting this idea, *NRAS* and *KRAS* mutations are among the most common genetic alterations in AML, with *NRAS* mutations present in approximately 10–15% of cases and *KRAS* mutations in about 5%^18^. RAS pathway activation is also a well-established mechanism of FLT3i resistance and is observed in approximately one-third of patients with *FLT3*-mutated AML who develop resistance to FLT3is^14,18^. Consistent with these previous clinical and laboratory findings, we confirmed that two *NRAS*-mutant AML models harboring G12C and Q61K mutations were resistant to gilteritinib and AC220 (Fig. 1B and data not shown). Based on these observations, we hypothesized that *SPRY* loss promotes FLT3i resistance by activating RAS signaling in AML. Indeed, *SPRY3* knockout induced RAS pathway activation in AML cells (Fig. 1C), consistent with its established role as an upstream negative regulator of RTK/RAS signaling.

**Figure 1.**
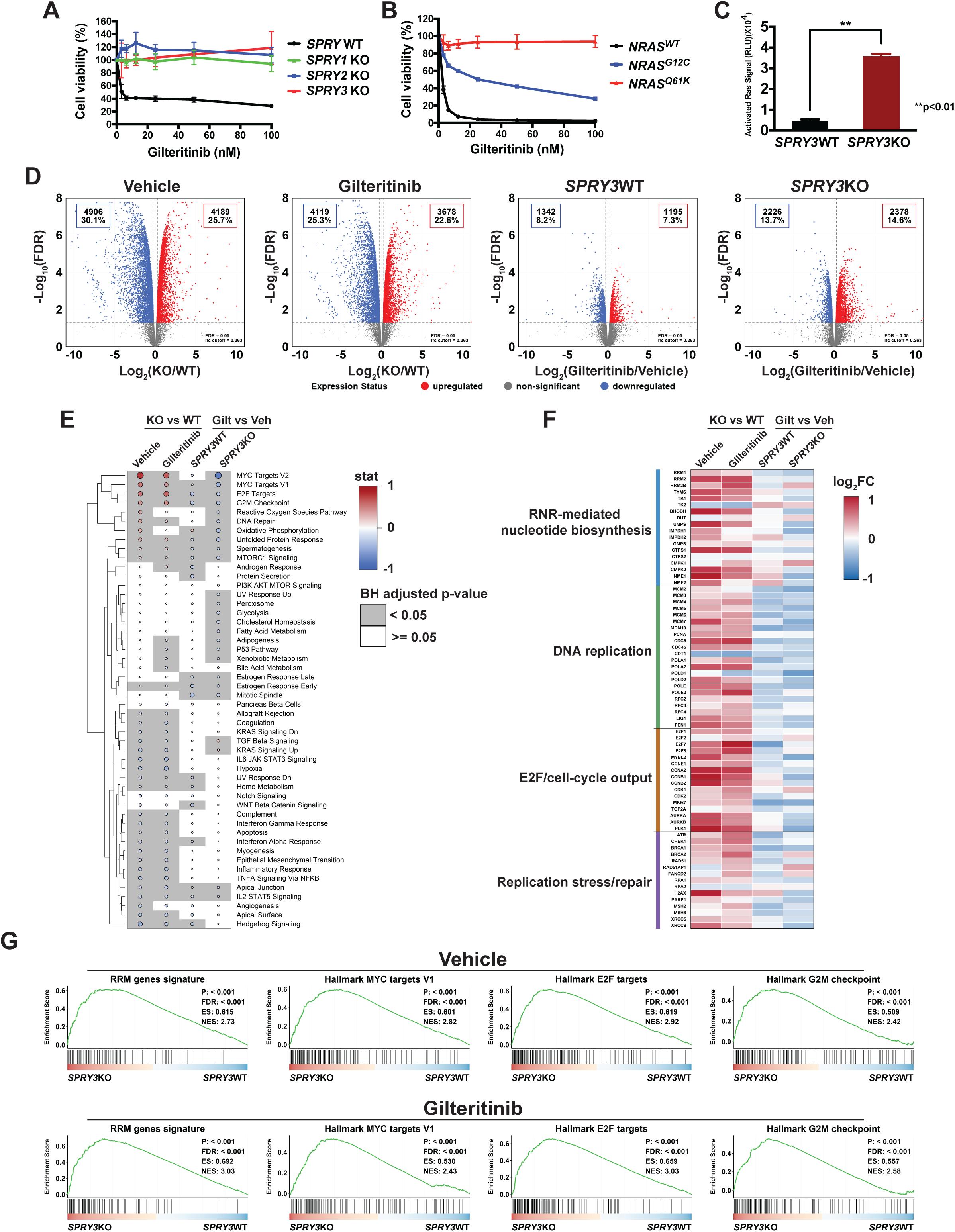
*SPRY3* loss drives gilteritinib resistance through RAS activation and MYC/E2F1-dependent RNR upregulation in *FLT3*^ITD^ AML. **A.** Growth curves of wild-type (WT) and *SPRY1*-, *SPRY2*-, or *SPRY3*-knockout (KO) *FLT3*^ITD^+ MV4-11 AML cells following treatment with gilteritinib. Loss of *SPRY1*, *SPRY2*, or *SPRY3* significantly increased cell viability in the presence of gilteritinib, indicating that knockout of *SPRY* family genes confers resistance to FLT3i in AML. **B.** Growth curves of *NRAS*^WT^/*FLT3*^ITD^+ and *NRAS*^mut^ MV4-11 AML cells following gilteritinib treatment. *NRAS*^mut^ cells showed significantly increased viability in the presence of gilteritinib, indicating that *NRAS* mutations confer resistance to FLT3i in AML. **C.** RAS pathway signaling in *FLT3*^ITD^+ MV4-11 AML cells was measured using a Ras GTPase Chemi ELISA kit, demonstrating that *SPRY3* loss leads to activation of RAS signaling. **D.** Volcano plots showing differential gene expression in *SPRY3*KO vs *SPRY3*WT cells under vehicle or gilteritinib treatment, and gilteritinib-induced transcriptional responses in *SPRY3*WT and *SPRY3*KO cells. **E.** Hallmark pathway activity dotplot showing pathway-level transcriptional changes across the same comparisons; dot color and size indicate pathway statistic, and gray shading indicates BH-adjusted P < 0.05. **F.** Heatmap of selected RNR-mediated nucleotide biosynthesis, DNA replication, E2F/cell-cycle, and replication stress/repair genes across the indicated comparisons. **G.** GSEA running enrichment plots for the *RRM2* 126-gene signature, *MYC* targets, *E2F* targets, and G2M checkpoint gene sets in *SPRY3*KO versus *SPRY3*WT cells under vehicle and gilteritinib treatment.

### *SPRY3* loss and *NRAS* active mutations promotes RNR upregulation upon FLT3i treatment in *FLT3*^ITD^+ AML

To investigate the transcriptional mechanisms by which SPRY3/RAS signaling promotes FLT3i resistance, we performed RNA-seq in *SPRY3*-deficient and WT AML cells treated with either vehicle or gilteritinib. Under vehicle conditions, *SPRY3* deficiency induced widespread transcriptional reprogramming, with approximately 30.1% and 25.7% of genes significantly downregulated and upregulated, respectively, compared with WT cells (Fig. 1D). A similarly extensive transcriptional difference was observed following gilteritinib treatment, indicating that *SPRY3* deficiency profoundly reshapes the AML transcriptome under both basal conditions and FLT3 inhibition. Hallmark gene set enrichment analysis (GSEA) demonstrated that *SPRY3* deficiency was associated with significant enrichment of MYC targets, E2F targets, G2/M checkpoint, DNA repair, and other proliferation-associated pathways under both vehicle- and gilteritinib-treated conditions (Fig. 1E), suggesting activation of a MYC/E2F-driven proliferative transcriptional program.

To identify the downstream effectors underlying these pathway alterations, we examined genes involved in nucleotide biosynthesis, DNA replication, cell-cycle progression, and replication stress responses (Fig. 1F). *SPRY3*-deficient cells consistently exhibited increased expression of RNR-mediated nucleotide biosynthesis genes, including *RRM1*, *RRM2*, *RRM2B*, *TYMS*, and *TK1*, together with genes involved in DNA replication (*MCM2–7*, *POLA1/2*, *POLD1/2*, *POLE*, *RFCs*, and *PCNA*), E2F-driven cell-cycle progression (*E2F1–3*, *CCNA2*, *CCNB1/2*, *CDK1*, *PLK1*, and *AURKA/B*), and replication stress responses (*CHEK1*, *BRCA1/2*, *RAD51*, *FANCD2*, *RPA1*, and *PARP1*). These data suggest that activation of RNR-mediated nucleotide biosynthesis represents a major downstream consequence of the MYC/E2F transcriptional program.

To further validate this model, we performed competitive gene set testing on curated RRM gene signatures^30,31^ and hallmark gene sets^32^. The RRM signature was significantly enriched in *SPRY3*-deficient cells under both vehicle- and gilteritinib-treated conditions and closely paralleled the enrichment of MYC targets, E2F targets, and G2/M checkpoint gene sets (Fig. 1G). These findings further support a model in which RNR-mediated nucleotide biosynthesis is an integral downstream component of the MYC/E2F-driven transcriptional network, providing the nucleotide supply required to sustain DNA replication and leukemic cell survival and proliferation during FLT3i treatment.

In parallel, a high-throughput small-molecule screen of 320 FDA-approved drugs identified clofarabine, an FDA-approved RNR inhibitor, as a strong hit that restored gilteritinib sensitivity in *SPRY3*KO/*FLT3*^ITD^+ AML cells, with the MAPK inhibitor trametinib used as a positive control. Interestingly, other RNR inhibitors, including cladribine and gemcitabine, were also identified in this drug screen (data not shown). Together, these complementary approaches converged on RNR activation as a key downstream mechanism of RAS-driven FLT3i resistance in AML.

### MAPK signaling and MYC/E2F1-dependent RNR induction mediate FLT3i resistance in NRASmut AML

We next validated this finding by real-time PCR and found that FLT3i treatment markedly upregulated all three RNR subunits, RRM1, RRM2, and RRM2B, in both *SPRY3*KO/*FLT3*^ITD^+ and *NRAS*-mutant (G12C and Q61K)/*FLT3*^ITD^+ AML cells (Supplementary Fig. S1B–E). Because *SPRY3*KO and *NRAS*-mutant AML cells are the primary models used in this study to investigate FLT3i resistance, the terms “*SPRY3*KO” and “*NRAS*^mut^” hereafter refer to *SPRY3*KO/*FLT3*^ITD^+ and *NRAS*^mut^/*FLT3*^ITD^+ AML cells, respectively. To determine whether this RNR induction depends on upstream RAS/MAPK signaling, we treated *NRAS*^mut^ AML cells with the MEK inhibitor trametinib. Trametinib markedly suppressed the gilteritinib-induced upregulation of *RRM1*, *RRM2*, and *RRM2B* (Fig. 2A–C), placing the RNR transcriptional response downstream of MAPK signaling. Clofarabine similarly reduced the gilteritinib-induced increase in all three RRM transcripts (Fig. 2A–C), indicating that pharmacologic RNR inhibition is accompanied by attenuation of this adaptive RNR expression program. Consistent with previous studies showing that E2F1 and MYC regulate RNR expression^33^, we next examined their contribution to gilteritinib-induced RRM transcription. *E2F1* knockdown strongly suppressed *RRM2* induction and also reduced the induction of RRM1 and RRM2B in *NRAS*^mut^ AML cells (Fig. 2D–F). *MYC* knockdown reduced gilteritinib-induced *RRM1* and *RRM2B* expression, while having a more limited effect on *RRM2* (Fig. 2G–I). Importantly, knockdown of either *E2F1* or *MYC* restored gilteritinib sensitivity in *NRAS*^mut^ AML cells (Fig. 2J, K), functionally linking the MYC/E2F1 transcriptional program to FLT3i resistance. To directly determine whether individual RNR subunits are required for the resistant phenotype, we used siRNAs to deplete *RRM1*, *RRM2*, or *RRM2B*. Silencing any of the three subunits significantly restored gilteritinib sensitivity in *NRAS*^mut^ AML cells (Fig. 2L–N). Together, these results support a model in which RAS/MAPK activation drives a MYC/E2F1-associated RNR transcriptional program that is required to sustain FLT3i resistance.

**Figure 2.**
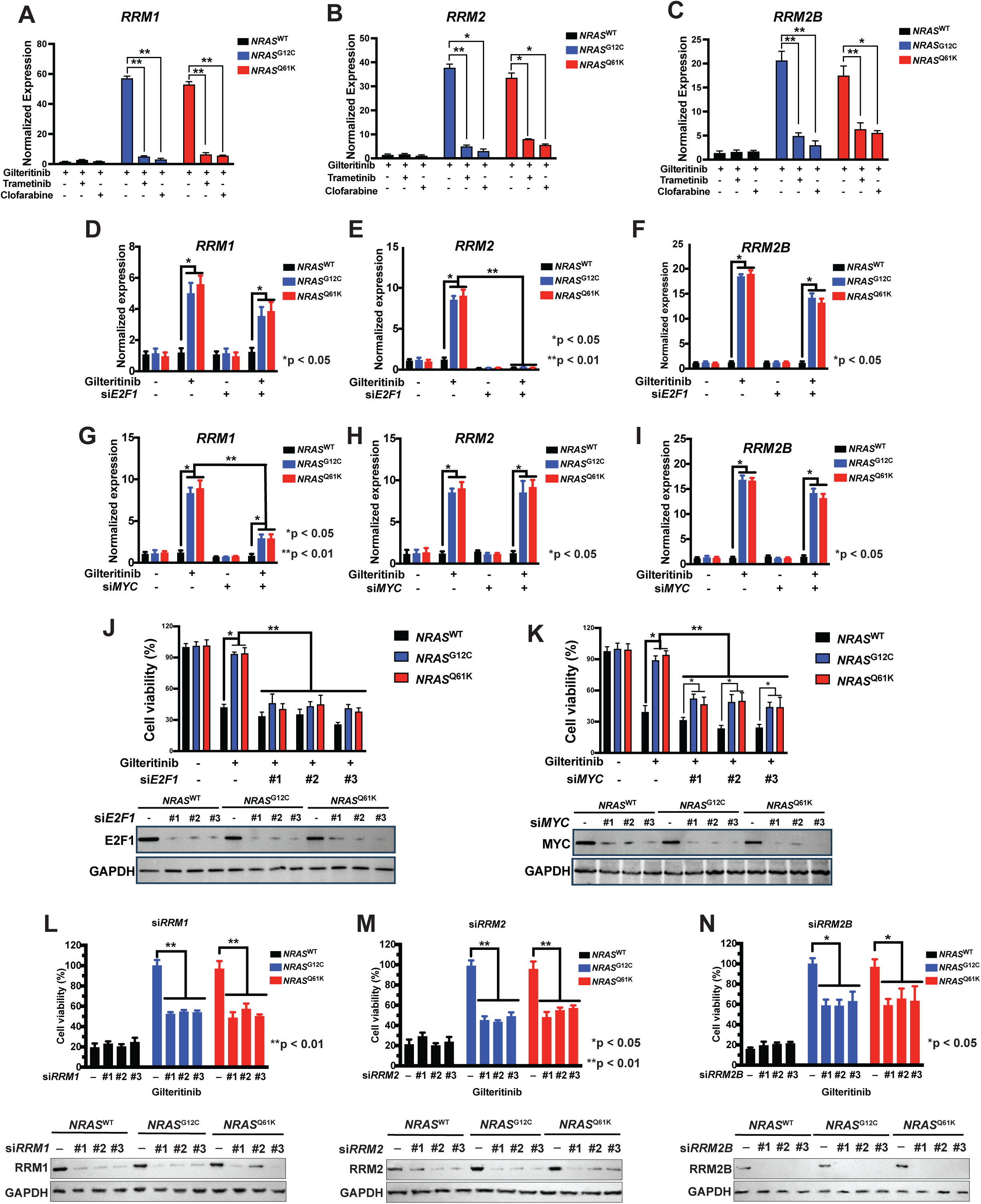
MAPK signaling and MYC/E2F1 drive gilteritinib-induced *RRM* upregulation, while depletion of RNR subunits restores gilteritinib sensitivity in *NRAS*^mut^/*FLT3*^ITD^+ AML cells. A-C. MV4-11 AML cells expressing *NRAS*^WT^, *NRAS*^G12C^, or *NRAS*^Q61K^ were treated with gilteritinib alone or together with trametinib or clofarabine. Expression of (A) *RRM1*, (B) *RRM2*, and (C) *RRM2B* was measured by RT-qPCR. D-F. Effects of *E2F1* knockdown on gilteritinib-induced expression of (D) *RRM1*, (E) *RRM2*, and (F) *RRM2B* in *NRAS*^WT^ and *NRAS*^mut^ MV4-11 AML cells. G-I. Effects of *MYC* knockdown on gilteritinib-induced expression of (G) *RRM1*, (H) *RRM2*, and (I) *RRM2B*. J. *E2F1* knockdown restored gilteritinib sensitivity in *NRAS*^mut^ AML cells; immunoblotting below confirms *E2F1* knockdown. K. *MYC* knockdown restored gilteritinib sensitivity in *NRAS*^mut^ AML cells; immunoblotting below confirms *MYC* knockdown. L-N. MV4-11 AML cells expressing *NRAS*^WT^, *NRAS*^G12C^, or *NRAS*^Q61K^ were transfected with three independent siRNAs targeting (L) *RRM1*, (M) *RRM2*, or (N) *RRM2B* and then treated with gilteritinib. Cell viability is shown above, and immunoblotting below confirms knockdown of the corresponding RNR subunit. Data are presented as mean ± SD from three replicates. Statistical significance is indicated in the figure.

### Pharmacologic RNR inhibition reverses FLT3i resistance in *SPRY3*-deficient and *NRAS*^mut^ *FLT3*^ITD^+ AML cells

Having established that RNR activation functionally contributes to FLT3i resistance, we next investigated whether pharmacologic inhibition of RNR could restore FLT3i sensitivity in *FLT3*^ITD^+ AML cells. To this end, we evaluated multiple direct and indirect RNR inhibitors (Fig. 3A). We first examined the well-characterized RNR inhibitor clofarabine and found that it effectively reversed resistance to both AC220 and gilteritinib in *SPRY3*KO and *NRAS*^mut^ AML cells in both MV4-11 and MOLM-14 cells, which harbor homozygous and heterozygous *FLT3*^ITD^, respectively (Fig. 3B–E). Notably, clofarabine alone exhibited minimal cytotoxicity at the concentrations tested (Fig. 3F), indicating that its effect primarily reflects restoration of FLT3i sensitivity rather than nonspecific cytotoxicity.

**Figure 3.**
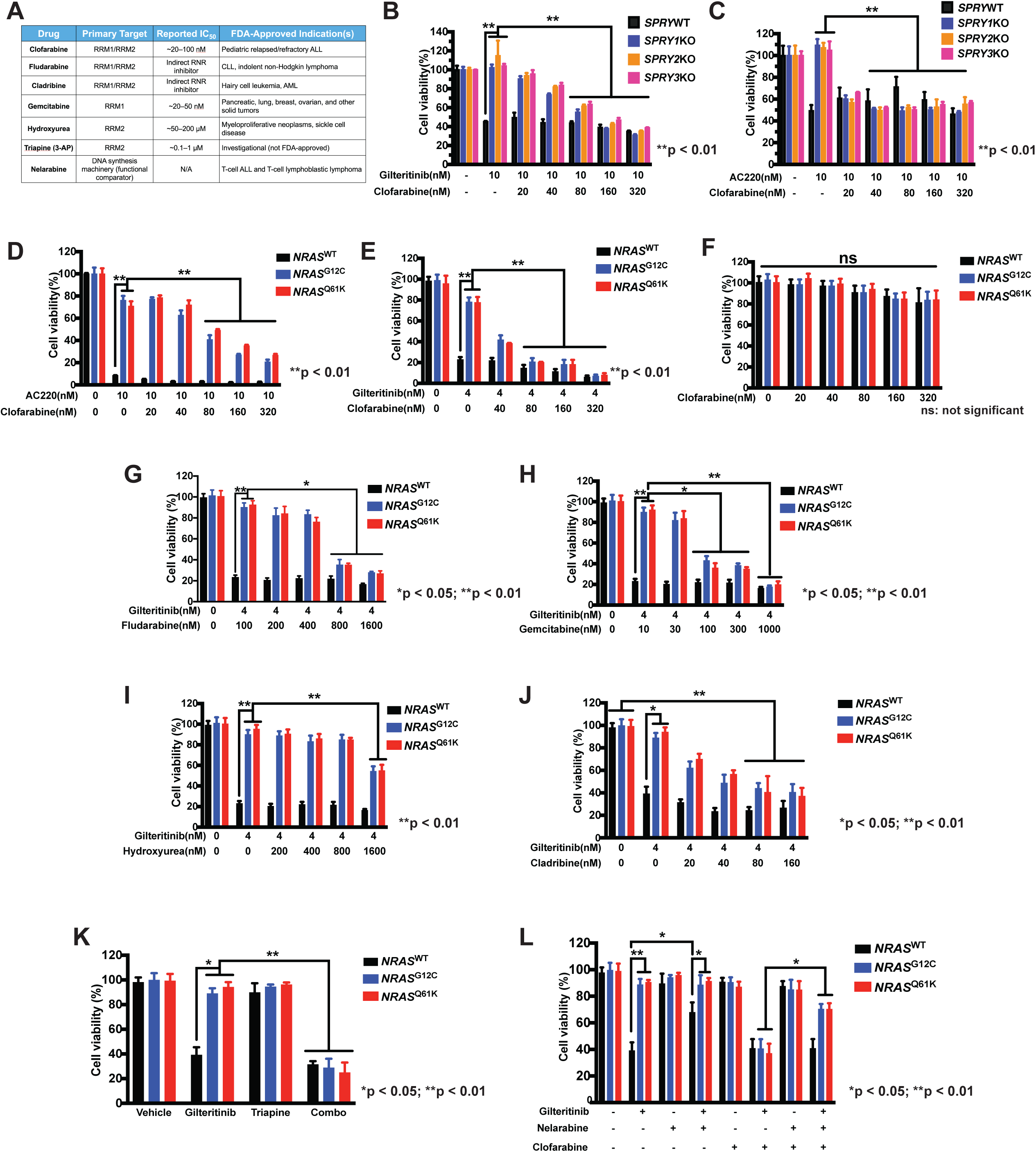
Pharmacologic RNR inhibition reverses FLT3i resistance in *SPRY3*KO/*FLT3*^ITD^+ and *NRAS*^mut^/*FLT3*^ITD^+ AML cells. **A-B**. Clofarabine treatment effectively reversed AC220 (A) and gilteritinib (B) resistance in *SPRY3*KO/*FLT3*^ITD^+ MV4-11 AML cells. **C-D.** Clofarabine treatment effectively reversed AC220 (C) and gilteritinib (D) resistance in *NRAS*^mut^/*FLT3*^ITD^+ MV4-11 AML cells. **E.** Clofarabine alone exhibited minimal cytotoxicity in MV4-11 AML cells. **F-H.** Additional RNRis, including fludarabine, gemcitabine, and hydroxyurea, also effectively reversed gilteritinib resistance in *NRAS*^mut^/*FLT3*^ITD^+ MV4-11 AML cells. **I**. Cladribine, a clinically approved purine nucleoside analog and indirect RNR inhibitor, reversed FLT3i resistance in *NRAS*^mut^/*FLT3*^ITD^+ AML cells. **J.** Triapine, a potent RNRi, reversed FLT3i resistance in *NRAS*^mut^/*FLT3*^ITD^+ MV4-11 AML cells. **K.** Nelarabine, an RNR activator, induced FLT3i resistance in *NRAS*^WT^ MV4-11 AML cells; this effect was reversed by clofarabine. Data represent the means ± SD of three replicates.

Consistent with these findings, additional RNR inhibitors, including fludarabine, gemcitabine, and hydroxyurea, also reversed FLT3i resistance in *NRAS*^mut^ AML cells in both homozygous and heterozygous *FLT3*ITD backgrounds (Fig. 3G–I and data not shown), while generally exhibiting minimal cytotoxicity as single agents (Supplementary Fig. 2A–C). We further examined cladribine, a clinically approved purine nucleoside analog and indirect RNR inhibitor^34^, and found that it similarly restored FLT3i sensitivity in *NRAS*^mut^ AML cells (Fig. 3J). Triapine, a potent RNR inhibitor that has been evaluated in multiple Phase I and II clinical trials for solid tumors and hematologic malignancies, including AML and CML^35^, also reversed FLT3i resistance in *NRAS*^mut^ AML cells while exhibiting minimal single-agent toxicity at the concentrations tested (Fig. 3K and data not shown). To further test the relationship between RNR activity and FLT3i resistance from the opposite direction, we examined nelarabine, a recently described RNR activator^31,36^. Nelarabine induced gilteritinib resistance in *NRAS*-wild-type AML cells, and this effect was reversed by clofarabine (Fig. 3L). Together, these complementary pharmacologic studies demonstrate that RNR activity is closely linked to FLT3i response and show that RNR inhibition consistently restores FLT3i sensitivity across multiple genetic models and pharmacologic approaches.

### The RNRi clofarabine disrupts *NRAS*-driven upregulation of nucleotide metabolism and DNA replication

Building on our identification of the RAS/RNR pathway as a mediator of FLT3i resistance in AML, we next sought to define the molecular mechanism by which clofarabine overcomes FLT3i resistance in *NRAS*^mut^ AML. We first examined the effects of gilteritinib alone or in combination with clofarabine on cell proliferation and apoptosis. Under gilteritinib treatment, *NRAS*^mut^ AML cells exhibited markedly increased proliferation, as measured by Ki-67 staining, and reduced apoptosis, as assessed by Annexin V staining, compared with *NRAS*^WT^ cells. Importantly, clofarabine significantly reversed this high-proliferation, low-apoptosis phenotype in *NRAS*^mut^ AML cells (Fig. 4A-B). Additionally, we examined whether *NRAS* mutations or RNR inhibition altered the cell-cycle response of *FLT3*^ITD^ AML cells to FLT3 inhibition. Gilteritinib increased the G1/G0 population and decreased the G2/M population in both *NRAS*^WT^ and *NRAS*^mut^ cells. Clofarabine alone had only modest effects on cell-cycle distribution, and the combination of clofarabine and gilteritinib did not produce substantial additional changes compared with either single agent (Supplementary Fig. 3A, B). These findings suggest that the enhanced therapeutic efficacy of combined RNR and FLT3 inhibition is not primarily mediated by changes in cell-cycle distribution.

**Figure 4.**
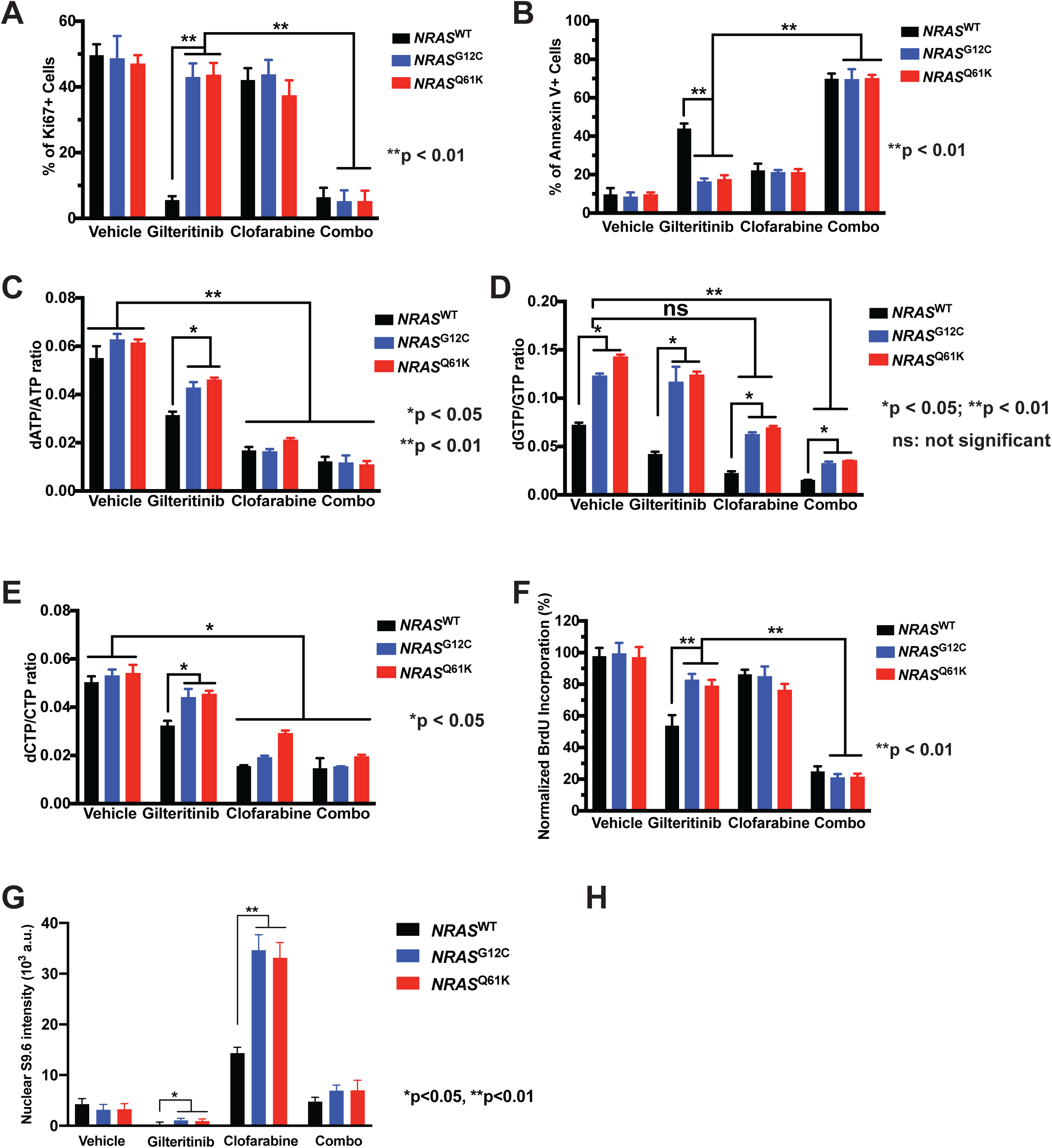
Clofarabine reverses gilteritinib-induced enhanced proliferation and reduced apoptosis in *NRAS*^mut^/*FLT3*^ITD^+ AML cells and disrupts NRAS-driven upregulation of nucleotide metabolism and DNA replication. MV4-11 AML cells harboring *NRAS*^WT^, *NRAS*^G12C^, or *NRAS*^Q61K^ mutations were treated with gilteritinib (10nM), clofarabine (40nM), or their combination. **A.** Proliferation was assessed by quantifying the percentage of Ki-67-positive cells, **B.** apoptosis was evaluated by measuring the percentage of Annexin V-positive cells. **C-E.** MV4-11 AML cells harboring *NRAS*^WT^, *NRAS*^G12C^, or *NRAS*^Q61K^ mutations were treated with gilteritinib (10nM), clofarabine (40nM), or their combination. Intracellular dNTP/NTP ratios, including (C) dATP/ATP, (D) dGTP/GTP, and (E) dCTP/CTP, were quantified by primer extension assay. **F.** DNA replication was assessed by BrdU incorporation following a 1-hour pulse prior to fixation and confocal imaging. Data represent the means ± SD of three replicates.

Because RNR plays a central role in maintaining nucleotide pool homeostasis and supplying deoxyribonucleotides for DNA replication^23^, we hypothesized that RNR activation in resistant cells promotes aberrant nucleotide accumulation to sustain replication. To test this hypothesis, we first measured dNTP/NTP levels using a primer extension–based assay^37^ in gilteritinib-treated *NRAS*^mut^ AML cells compared with *NRAS*^WT^ AML cells and found that gilteritinib treatment caused an uneven increase in dNTP pools. This imbalance was characterized by a marked elevation of dGTP levels, whereas clofarabine treatment effectively reversed this increase (Fig. 4C-E), suggesting that clofarabine restores nucleotide pool homeostasis in FLT3i-resistant AML cells.

Given that RNR is essential for DNA replication^22–24^, we next assessed replication dynamics using a BrdU incorporation assay. We observed significantly increased DNA replication in gilteritinib-treated *NRAS*^mut^ AML cells compared with *NRAS*^WT^ AML cells. Remarkably, the addition of clofarabine significantly reversed the increased replication observed in *NRAS*^mut^ AML cells (Fig. 4F), suggesting that enhanced DNA replication is a key downstream consequence of RNR activation in this context. To further investigate the underlying molecular mechanism, we performed RNA-seq analysis of *NRAS*^G12C^ and *NRAS*^WT^ AML cells treated with vehicle, gilteritinib, clofarabine, or the combination of gilteritinib and clofarabine (Supplementary Fig. 4). Differential gene expression analysis showed that each treatment induced widespread transcriptional changes in *NRAS*^G12C^ AML cells compared with *NRAS*^WT^ cells, with gilteritinib alone and the combination treatment producing the most extensive transcriptional reprogramming (Supplementary Fig. 4A). Gene-level analysis further demonstrated that *NRAS*^G12C^ cells maintained elevated expression of genes involved in RNR-mediated nucleotide biosynthesis, DNA replication, cell-cycle progression, and DNA damage/repair following gilteritinib treatment, consistent with persistent activation of a high-replication transcriptional program despite FLT3 inhibition (Supplementary Fig. 4B). Consistent with these findings, Hallmark pathway analysis demonstrated persistent enrichment of MYC targets, E2F targets, and other proliferation-associated pathways in *NRAS*^G12C^ cells (Supplementary Fig. 4C). Furthermore, GSEA using a curated RRM2 126-gene signature showed that the RRM2-associated transcriptional program closely paralleled the enrichment of MYC and E2F target gene sets across all treatment groups (Supplementary Fig. 4D), supporting a close functional relationship between MYC/E2F signaling and RNR-mediated nucleotide biosynthesis. Together with our functional studies, these findings support a model in which oncogenic *NRAS* creates a dependence on RNR-mediated nucleotide biosynthesis to sustain DNA replication under FLT3i treatment. Clofarabine overcomes FLT3i resistance by disrupting this metabolic dependency, leading to dNTP depletion, excessive replication stress, and apoptotic cell death.

### RNR inhibition overcomes NRAS-driven FLT3i resistance in AML cell line–derived xenograft (CDX) models *in vivo*

To further establish the clinical relevance of our findings, we used two complementary models of RAS pathway activation: *SPRY3* deficiency, as a genetically defined model, and *NRAS* mutation, as a clinically relevant driver of FLT3i resistance. We first evaluated the therapeutic efficacy of combined FLT3 and RNR inhibition in *SPRY3*KO AML using cell line-derived xenograft (CDX) models, with *SPRY3*^WT^/*FLT3*^ITD^+ cells serving as controls. AML cells were transduced with a firefly luciferase-expressing lentiviral construct and injected into NSG (Nod.Cg-Prkdc^scid^Il2rg^tm1Wjl^/SzJ) immunocompromised mice. After engraftment of AML was confirmed, mice were randomized into four treatment groups: vehicle, gilteritinib (30 mg/kg, po, qod), clofarabine (40 mg/kg, ip, qod), or the combination. Treatments were administered three times per week for a total of four doses, and leukemic burden was monitored by *in vivo* bioluminescence imaging (BLI) after 12 days (Supplementary Fig. 5A).

In the *SPRY3*WT model, both gilteritinib monotherapy and the combination treatment effectively reduced leukemic burden and prolonged survival, whereas clofarabine alone had no significant effect compared with vehicle (Supplementary Fig. 5B–C). In contrast, in the *SPRY3*KO FLT3i-resistant CDX model, gilteritinib monotherapy showed minimal efficacy, consistent with marked gilteritinib resistance, and clofarabine alone exerted only limited antileukemic activity (Supplementary Fig. 5B and F). Notably, the combination of gilteritinib and clofarabine significantly reduced leukemic burden and prolonged survival in *SPRY3*KO recipient mice (Supplementary Fig. 5D and G). No treatment-related weight loss was observed, supporting the tolerability of both agents (Supplementary Fig. 5E and H). Together, these findings demonstrate that RNR inhibition effectively overcomes gilteritinib resistance driven by *SPRY3* loss *in vivo*.

To determine whether these findings extend to clinically relevant *NRAS*-driven resistance, we next evaluated the same therapeutic strategy in *NRAS*^mut^ AML CDX models. Consistent with prior reports and our *in vitro* data, *NRAS*^G12C^ model exhibited marked resistance to gilteritinib treatment (30 mg/kg/day, po, qod for 4 times) compared to *NRAS*^WT^/*FLT3*^ITD^+ controls *in vivo*. Importantly, treatment with the clinically used RNRi clofarabine (40 mg/kg, ip, qod for 4 times) effectively re-sensitized *NRAS*^G12C^ AML cells to gilteritinib *in vivo*, resulting in a significant reduction in leukemic burden and prolonged survival (Fig. 5A, B, C, D, F, G). Clofarabine monotherapy showed only modest effects, and no treatment-related weight loss was observed in any group (Fig. 5E, H), indicating minimal toxicity of both drugs.

**Figure 5.**
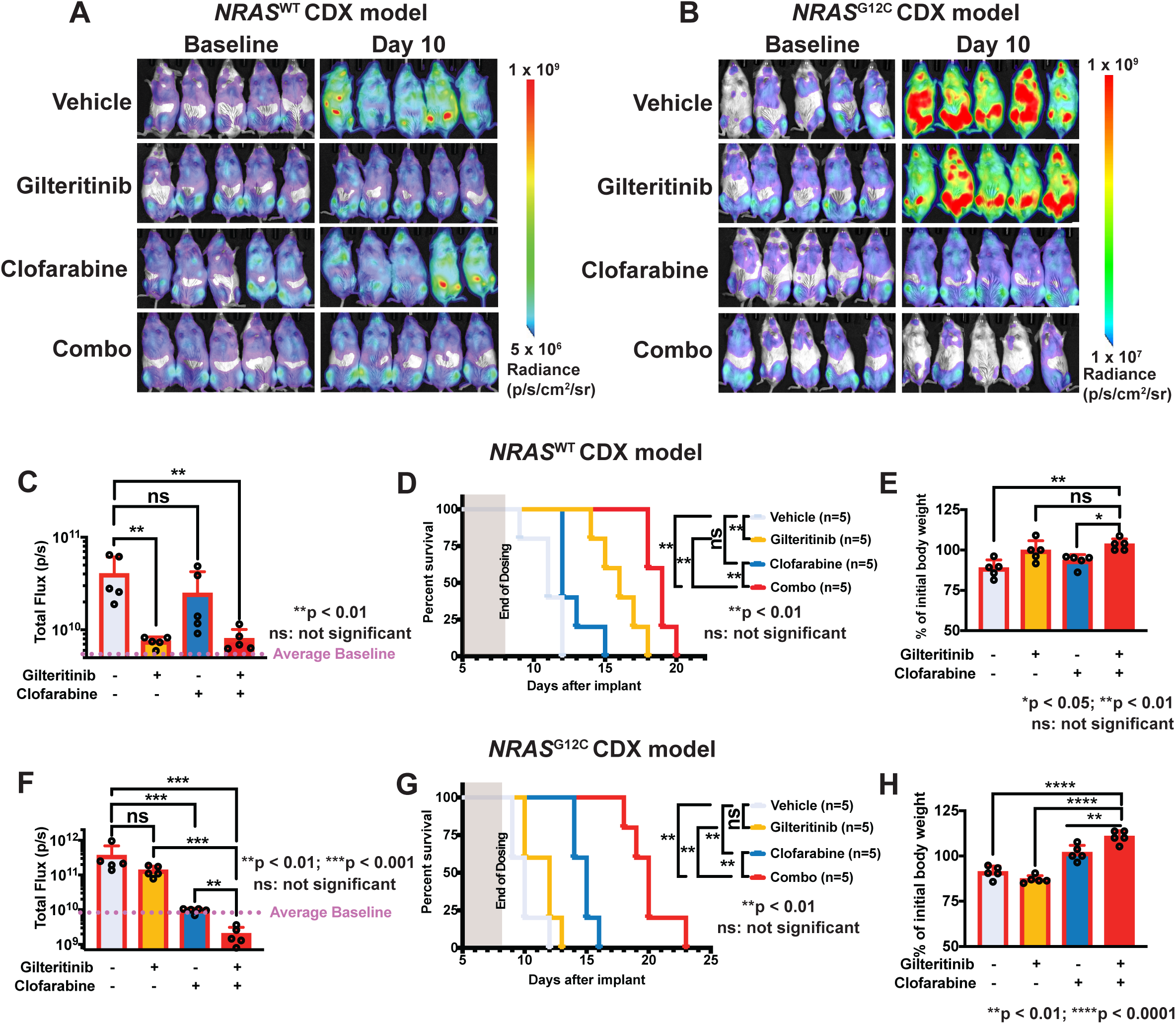
Clofarabine reverses *NRAS*^G12C^-induced gilteritinib resistance in CDX models of AML. **A.** Images of *in vivo* BLI assessment of NSG mice engrafted with *NRAS*^WT^ and *NRAS*^G12C^ luciferase-tagged MOLM-14 cells during the experiment. **B and E.** Quantification of BLI data from the (B) *NRAS*^WT^ and (E) *NRAS*^G12C^ CDX studies. C and F. Kaplan-Meier curves representing survival data of subjects from each treatment group in (C) *NRAS*^WT^ and (F) *NRAS*^G12C^ CDX studies. D and G. The combination treatment had no effect on body weight of recipients from both (D) *NRAS*^WT^ and (G) *NRAS*^G12C^ CDX studies. Data represent the means ± SD (n = 5/group).

Collectively, these results demonstrate that combined FLT3 and RNR inhibition represents an effective therapeutic strategy to overcome FLT3i resistance driven by NRAS pathway activation, whether through *SPRY3* loss or oncogenic *NRAS* mutation, in AML.

### RNR inhibition overcomes *NRAS*-driven FLT3i resistance in an *NRAS*^Q61K^/*FLT3*^ITD^+ patient-derived xenografts (PDX) model of AML

Because PDX models is a gold-standard platform in AML for evaluating therapeutic efficacy, we next sought to determine whether clofarabine re-sensitize *NRAS*^mut^ AML patient cells to gilteritinib with PDX models. To this end, we used a paired PDX model established from the same *FLT3*^ITD^+ AML patient, who was enrolled in a clinical trial performed in the mid-2010s of single-agent gilteritinib for relapsed and/or refractory *FLT3*-mutated AML (NCT02014558, NCT02421939) conducted at UCSF, the University of Pennsylvania, and Roswell Park Comprehensive Cancer Center. All patients received gilteritinib at FLT3-inhibitory doses (≥80 mg/day) and consented to institutional tissue banking protocols^38^. Samples were collected from this patient at diagnosis, when the leukemia was sensitive to gilteritinib, and again after relapse on gilteritinib treatment, when the disease had become resistant^38^. Importantly, RNA sequencing identified an *NRAS*^Q61K^ mutation at the time of acquired gilteritinib resistance^14^. For our *in vivo* studies, both the FLT3i-sensitive and FLT3i-resistant AML samples were successfully engrafted into NSG immunocompromised mice. Leukemia progression was monitored by flow-cytometric quantification of human CD45/CD33-expressing (hCD45+/hCD33+) cells in peripheral blood. Once engraftment was established after 60-75 days for gilteritinib-sensitive sample and 14-17 days for gilteritinib-resisatnt sample, recipients were randomized into four treatment groups (n = 5 per group): vehicle, gilteritinib (30 mg/kg, po, qod), clofarabine (40 mg/kg, ip, qod), or the combination. After six administrations, or when mice reached the predefined humane endpoint, animals were euthanized, and leukemic burden in bone marrow (BM), spleen, and peripheral blood (PB) was quantified by flow-cytometric analysis of hCD45+/hCD33+ cells (Fig. 6A).

**Figure 6.**
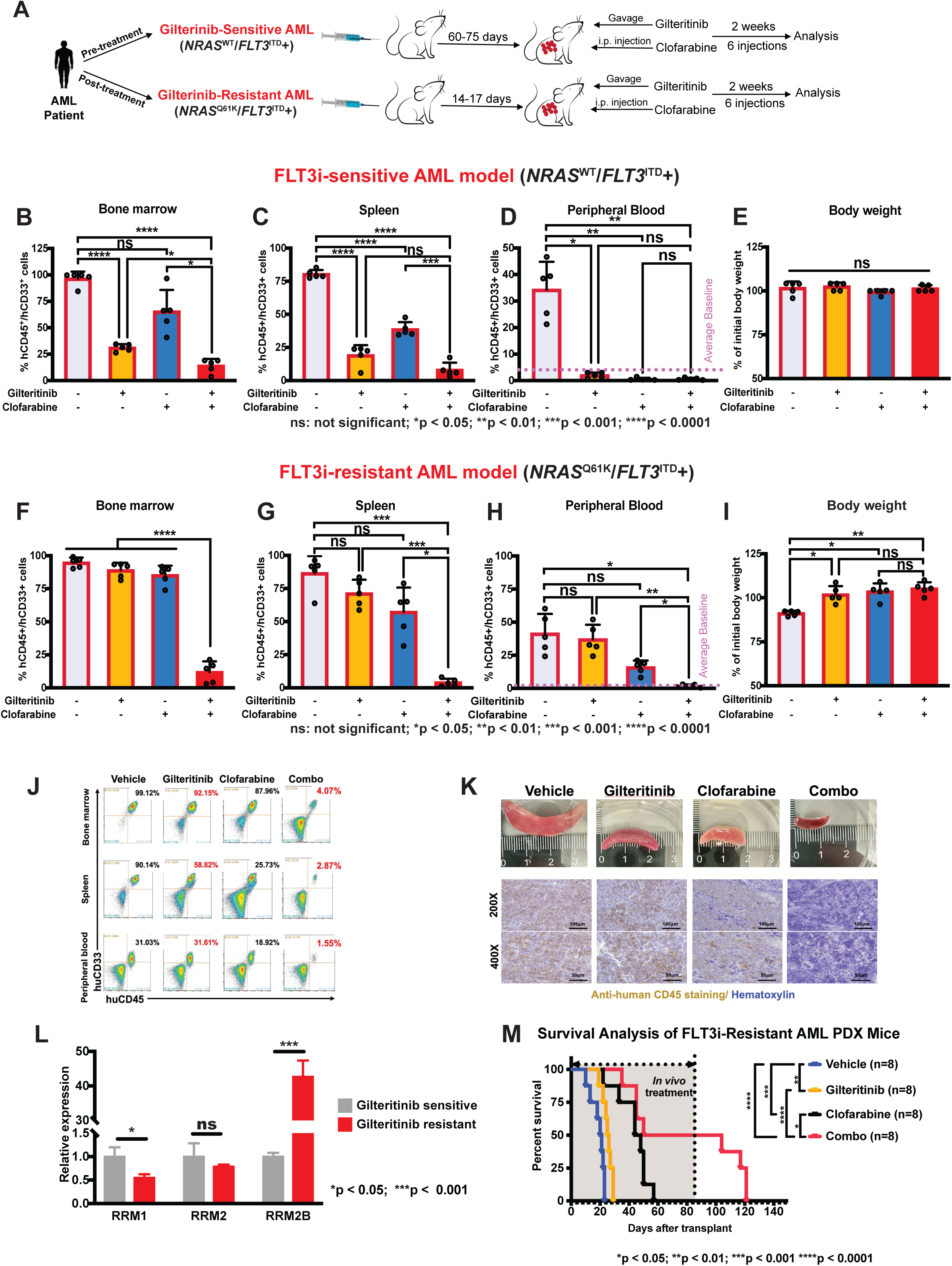
Clofarabine effectively reverses gilteritinib resistance in *NRAS*^Q61K^/*FLT3*^ITD^+ PDX models of AML and extends survival of AML recipients. **A.** Schematic representation of the PDX *in vivo* study design. **B-D.** Quantification of hCD45^+^ cells in the (B) bone marrow, (C) spleen, and (D) peripheral blood of the gilteritinib-sensitive model across the four treatment groups. **E.** No significant changes in recipient body weight were observed following combination treatment in the gilteritinib-sensitive model. **F-H.** Quantification of hCD45^+^ cells in the (F) bone marrow, (G) spleen, and (H) peripheral blood of the gilteritinib-resistant model across the four treatment groups. **I.** No significant changes in recipients’ body weight were observed following combination treatment in the gilteritinib-sensitive model; however, the vehicle group exhibited significant weight loss due to leukemia burden. **J.** Representative flow cytometry plots of human CD45^+^/CD33^+^(myeloid) cells in the bone marrow, spleen and peripheral blood of the gilteritinib-resistant PDX model, showing reduced leukemic burden following combination therapy. **K.** Representative images of spleen (top panel) and spleen sections and corresponding immunohistochemical staining for hCD45^+^ from the four treatment groups (bottom panel) in the gilteritinib-resistant model; Scale bars, 100 µm and 50 µm. **L.** RT-qPCR of human CD45^+^/CD33^+^ myeloid cells from the bone marrow of the gilteritinib-sensitive and gilteritinib-resistant models showing increased *RRM2B* expression in the gilteritinib-resistant model. **M.** Kaplan-Meier survival curves for each treatment group demonstrate that combination treatment significantly prolongs survival. Data represent the means ± SD (n = 8/group).

In the gilteritinib-sensitive PDX model, AML remained sensitive to gilteritinib monotherapy, with both gilteritinib alone and the combination treatment significantly reducing leukemic burden across BM, spleen and PB. Clofarabine monotherapy had minimal effects in the BM but resulted in a modest reduction of leukemic cells in the spleen (Fig. 6B and C and Supplementary Fig. 6A). Leukemic cells remained low in the PB across all treatment groups, likely reflecting slower disease progression in the FLT3i-sensitive model, which may limit systemic dissemination within the study timeframe (Fig. 6D). Consistent with these findings, IHC analysis showed a decrease in hCD45^+^ cells in the spleens of mice treated with either single agents or the combination (Supplementary Fig. 6B).

In contrast, in the FLT3i-resistant PDX model derived from the relapsed sample harboring an *NRAS*^Q61K^ mutation, gilteritinib monotherapy failed to reduce leukemic burden, confirming robust NRAS-driven FLT3i resistance *in vivo*. Importantly, combination treatment with gilteritinib and clofarabine produced a profound reduction in leukemic burden across BM, spleen, and PB, whereas clofarabine monotherapy had only minimal effects (Fig. 6F–H and Supplementary Fig. 6C). No significant body weight loss was observed in recipients receiving either monotherapy or combination treatment for 12 days, indicating good tolerability of both drugs. Body weight loss was observed only in the vehicle group, consistent with rapid disease progression in this cohort (Fig. 6I). Representative flow cytometry data of AML cells in the BM, spleen, and PB are shown in Fig. 7J. Consistent with the splenomegaly commonly observed in AML patients, the vehicle group exhibited marked splenomegaly, whereas combination treatment significantly reduced spleen size and weight (Fig. 6K top panel). Immunohistochemical (IHC) staining of spleen sections further showed a marked decrease in hCD45+ leukemic infiltration in mice receiving the combination treatment compared with those receiving single-agent therapy (Fig. 6K bottom panel). Strikingly, the expression of *RRM2B* was markedly upregulated in the gilteritinib-resistant AML sample compared with the gilteritinib-sensitive AML (Fig. 6L).

**Figure 7.**
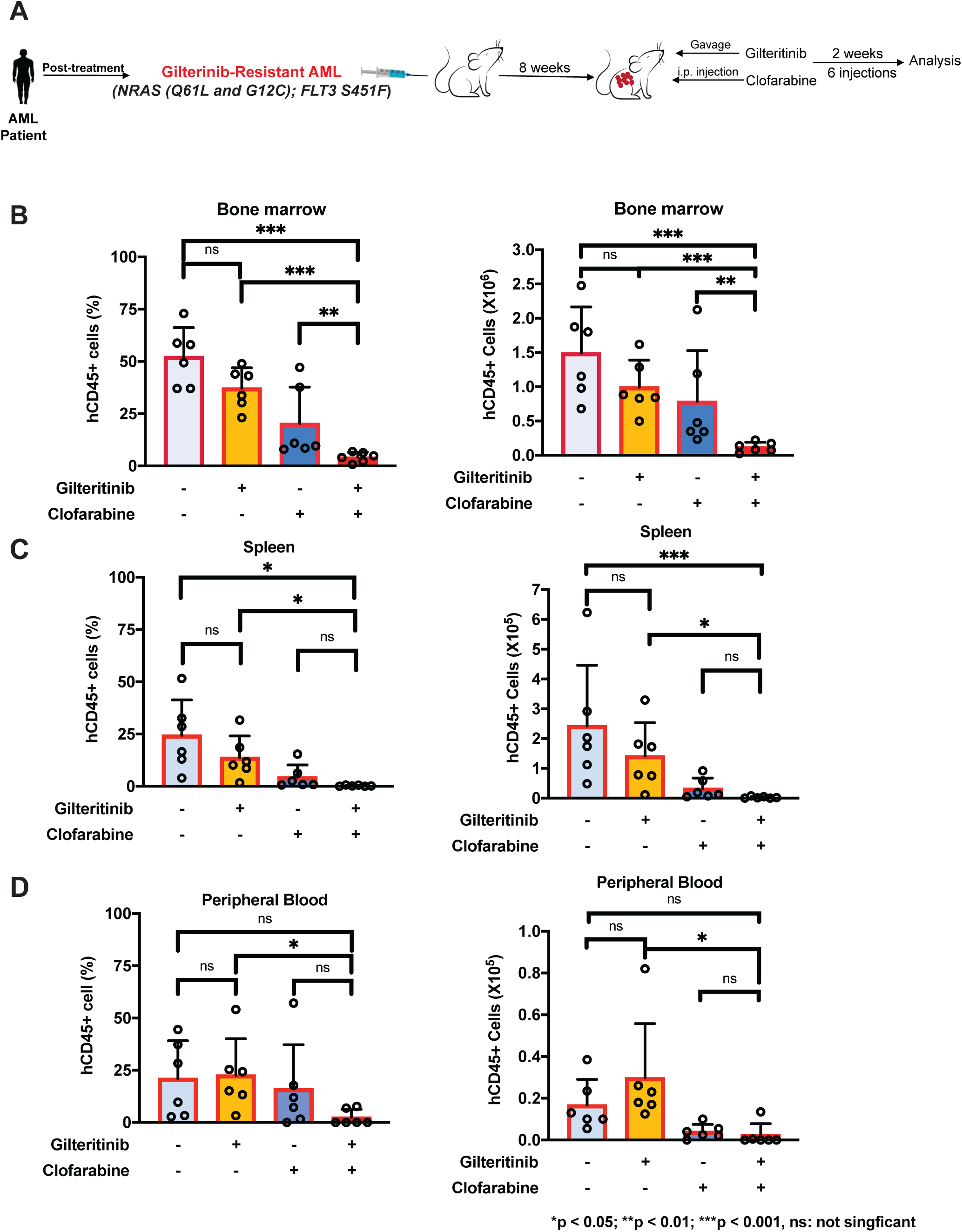
Clofarabine effectively reverses gilteritinib resistance in an *NRAS*^mut^/*FLT3* S451F AML PDX model. **A.** Schematic representation of the 2^nd^ PDX *in vivo* study design. **B–D.** NSG mice engrafted with a primary AML patient-derived xenograft (PDX) harboring *NRAS* (Q61L and G12C) and *FLT3 S451F* mutations were treated with vehicle, gilteritinib, clofarabine, or the combination of gilteritinib and clofarabine (n = 6 mice per group) for 2 weeks (6 total treatments). Leukemia burden was evaluated at the experimental endpoint by flow cytometric analysis of human CD45-positive (hCD45+) cells in the bone marrow (B), spleen (C), and peripheral blood (D). Both the percentage (left panels) and the absolute number (right panels) of hCD45+ cells are shown. Absolute bone marrow hCD45+ cell numbers were quantified from all four hind limb bones using counting beads. Each dot represents an individual mouse, and bars indicate the mean ± SD. The combination treatment significantly reduced leukemia burden compared with either gilteritinib or clofarabine monotherapy, particularly in the bone marrow, demonstrating that clofarabine effectively overcomes gilteritinib resistance in this AML PDX model.

**Figure 8.**
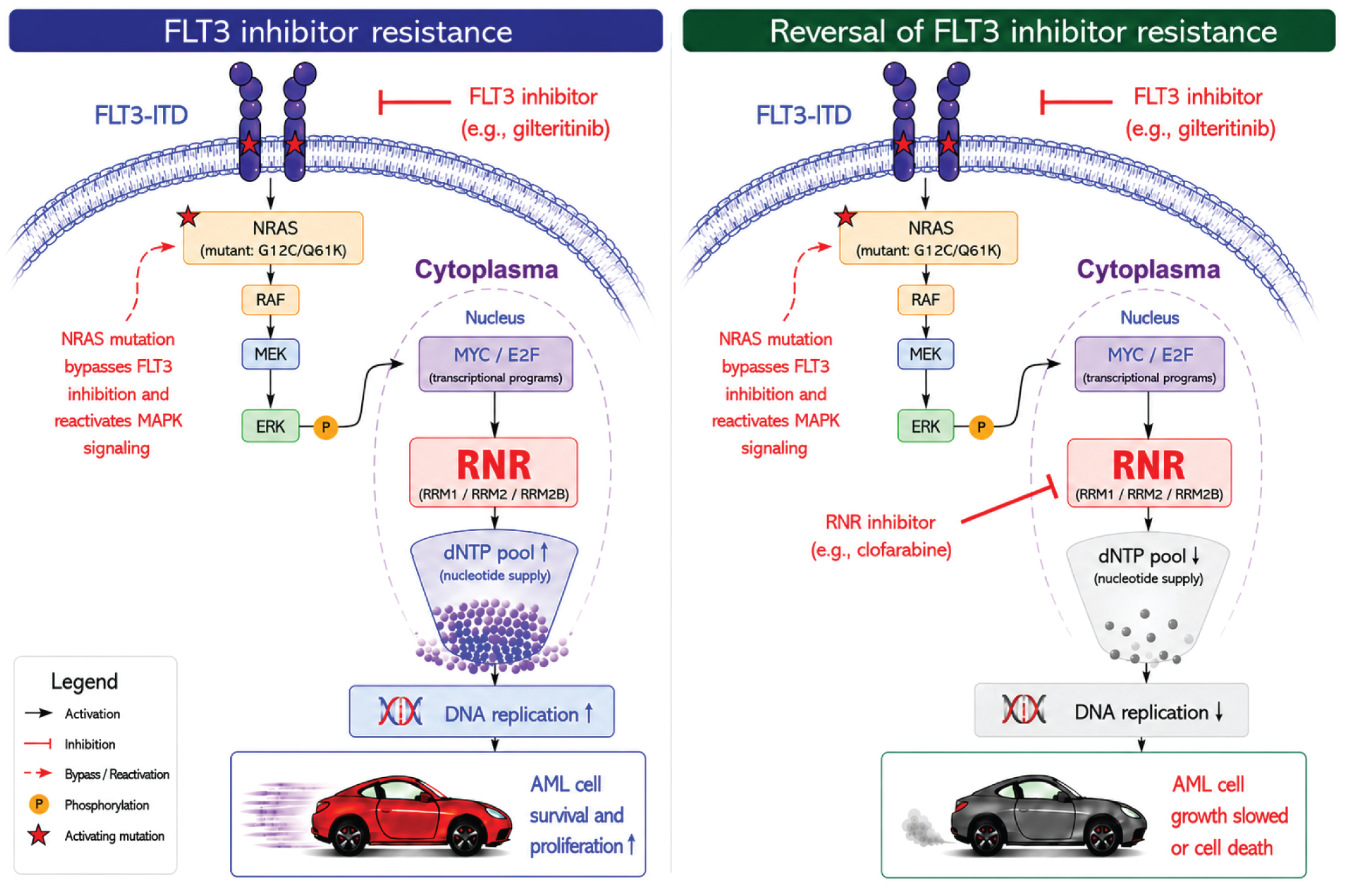
Schematic model showing that RNR inhibition reverses *NRAS*mut-mediated FLT3i resistance.

To further evaluate long-term therapeutic benefit, we performed an extended treatment study comparing gilteritinib monotherapy with combination therapy in the gilteritinib-resistant PDX model (n = 8 per group). All mice treated with gilteritinib monotherapy succumbed to disease within 40 days. In contrast, 4 of 8 mice receiving combination therapy remained alive at the end of the 12-week treatment period, and 2 survived beyond 120 days, including 40 days after treatment discontinuation, demonstrating a substantial survival advantage (Fig. 6M).

### RNR inhibition overcomes *NRAS*-driven FLT3i resistance in an *NRAS*^G12C^/*NRAS*^Q61L^/*FLT3*^S451F^ patient-derived xenografts (PDX) model of AML

To further validate that the RNRi clofarabine effectively overcomes *NRAS*-driven FLT3i resistance in primary AML, we identified an independent AML sample from the UPenn Xenograft Core harboring *STAG2* (p.E342Vfs*3), *NRAS* (Q61L and G12C), and *FLT3* S451F mutations. *FLT3* S451F is a rare, noncanonical *FLT3* variant that has been reported to confer constitutive kinase activation and limited sensitivity to FLT3is^39^. This genetically distinct AML PDX model, which carries *NRAS* (Q61L and G12C) mutations together with the *FLT3* S451F variant, provided an opportunity to evaluate the efficacy of the gilteritinib/clofarabine combination in an independent genetic background. NSG mice were engrafted with this AML sample, engraftment was confirmed at 6 weeks, and treatment was started at approximately 8 weeks after transplantation, as described above. Consistent with our previous findings, this AML PDX model exhibited limited sensitivity to gilteritinib monotherapy. In contrast, the combination of gilteritinib and clofarabine markedly reduced the leukemia burden compared with either gilteritinib or clofarabine alone, particularly in the bone marrow (Fig. 7A–D and Supplementary Fig. 7A-D). These findings further demonstrate that RNR inhibition effectively overcomes FLT3i resistance in genetically distinct *NRAS*-mutant AML PDX models.

Collectively, these results demonstrate that the RNRi clofarabine effectively overcomes *NRAS*-driven FLT3i resistance in primary AML samples. The combination of FLT3i and RNRi significantly reduces leukemic burden and prolongs survival of recipients, showing the strong translational potential of this therapeutic strategy for relapsed FLT3i-resistant *FLT3*^mut^+ AML.

## DISCUSSION

Acquired resistance to FLT3i remains a major obstacle to durable responses in FLT3-mutant AML^40,41^. Although reactivation of RAS/MAPK signaling is a well-established driver of resistance, how oncogenic signaling is translated into downstream programs that sustain leukemic survival under therapeutic pressure has remained poorly defined. Here, we identify ribonucleotide reductase (RNR)-dependent nucleotide metabolism as a critical and targetable effector of *NRAS*-driven FLT3i resistance. Our data support a model in which oncogenic RAS signaling enforces a metabolic state that sustains DNA replication in the presence of FLT3 inhibition. Activation of RAS signaling, through *SPRY3* loss or *NRAS* mutation, drives transcriptional upregulation of RNR subunits and expansion of intracellular dNTP pools, thereby maintaining nucleotide supply and enabling continued proliferation. This adaptive program is mediated, at least in part, through transcriptional regulators such as E2F and MYC, linking oncogenic signaling to nucleotide biosynthesis. Importantly, RNR activation is not simply a byproduct of proliferation but a functional requirement for resistance. Disruption of this pathway through pharmacologic or genetic inhibition of RNR reduces nucleotide synthesis, impairs DNA replication, and restores apoptotic responses in FLT3i-resistant AML cells. These effects are recapitulated *in vivo*, where combined FLT3 and RNR inhibition produces robust antileukemic activity in both CDX and PDX models of FLT3i-resistant AML.

These findings connect two previously distinct features of AML biology—RAS pathway activation and nucleotide metabolism—into a unified mechanism of therapeutic resistance. Although RAS/MAPK reactivation has been extensively implicated in relapsed AML^14,18,42^, and RNR activity has been broadly associated with cellular proliferation and drug resistance^25,27^, a direct functional link between these processes has not previously been established. Notably, prior studies have suggested that cancer cells can undergo adaptive RNR complex switching by dynamically altering RNR subunit composition to sustain deoxyribonucleotide production under therapeutic stress^27,43^. Consistent with this concept, we observed in our PDX model that RRM2B was the only RNR subunit markedly upregulated in resistant leukemia cells, suggesting selective RNR remodeling during FLT3i treatment. Our findings therefore support a model in which RAS pathway activation promotes adaptive RNR switching to maintain DNA replication and leukemia cell survival under FLT3 inhibition. By identifying RNR remodeling as a downstream consequence of RAS signaling, our study expands current paradigms of FLT3i resistance to include metabolic reprogramming of nucleotide synthesis as a central adaptive mechanism.

From a therapeutic perspective, our findings nominate RNR as a clinically actionable vulnerability in FLT3i-resistant AML. Notably, RNRis such as clofarabine and fludarabine are already FDA-approved, providing a clear path toward clinical translation. The consistent efficacy of combined FLT3 and RNR inhibition across multiple preclinical models, including PDX with paired diagnosis-relapse samples, indicates the potential of this strategy, particularly in the setting of RAS pathway activation. More broadly, these results support a new paradigm in which targeting metabolic dependencies downstream of oncogenic signaling can effectively overcome resistance to targeted therapies. Indeed, we found that clofarabine also sensitizes *NRAS*^mut^ AML cells to venetoclax (data not shown), suggesting that RNR inhibition may further enhance the efficacy of targeted therapies in *NRAS*^mut^ AML.

Clofarabine is a highly potent RNR inhibitor but is also associated with substantial toxicity, whereas cladribine is generally considered less toxic^44^. Although we observed minimal toxicity in both CDX and PDX models, we are now collaborating with a group at Cooper Hospital in New Jersey to initiate a new clinical trial repurposing cladribine to inhibit RNR and sensitize *NRAS*^mut^ AML to FLT3is.

The link between RNR and AML cell biology, as well as drug resistance, is increasingly appreciated. A previous study showed that RNR activation promotes AML differentiation^31^, which may initially seem inconsistent with its role in FLT3i resistance. However, RNR-driven differentiation occurs predominantly along the monocytic lineage^31^. Notably, recent studies have demonstrated that RAS pathway activation drives clonal selection and monocytic differentiation, thereby contributing to resistance to both FLT3is and venetoclax plus azacitidine^42,45^. Together with our findings, these observations suggest that therapy-induced RNR activation may promote monocytic differentiation as one mechanism of drug resistance in AML. In addition, RNR activation may contribute to resistance by promoting DNA replication, thereby facilitating gene amplification of resistance-associated oncogenes such as MYC or RAS. Gene amplification is a well-established mechanism of drug resistance in cancer^46^. Consistent with this possibility, RNR activation has been linked to KRAS inhibitor resistance in *KRAS*-mutant colorectal cancer (personal communication with Dr. Yaeger, Memorial Sloan Kettering Cancer Center). Another, non-mutually exclusive mechanism involves replication stress adaptation. Oncogenic *NRAS* induces replication stress (RS), characterized by dysregulated DNA replication and replication fork instability^47,48^. Persistent RS may render AML cells dependent on RNR-mediated dNTP synthesis to sustain DNA replication and survive FLT3 inhibition. RNR inhibition disrupts this adaptation, exacerbating replication stress beyond the cellular tolerance threshold and triggering replication catastrophe in FLT3i-resistant AML cells. Future studies will determine whether RNR activation contributes to lineage plasticity, gene amplification, replication stress adaptation, or other adaptive mechanisms that promote acquired FLT3i resistance in AML.

In summary, we identified RNR as a key vulnerability in *NRAS*-driven, FLT3i-resistant AML and defined a previously unrecognized RAS–RNR axis that enables leukemic cells to maintain nucleotide pools and sustain DNA replication under FLT3 inhibition. Our findings establish metabolic reprogramming of nucleotide synthesis as a core mechanism of FLT3i resistance and identify RNR inhibition as a tractable strategy to restore drug sensitivity. Together, these results provide a conceptual and translational framework for targeting oncogenic signaling–driven metabolic adaptations in AML and potentially other *RAS*-mutated malignancies.

## Authors’ Contributions

**Conception and design:** J. Huang

**Development of methodology:** Z. Tian, H. Zhang, J. Huang

**Acquisition of data (provided animals, acquired and managed patients, provided facilities, etc.):** Z. Tian, H. Zhang, S. Chatla, XY. Wei, Y. Liu, D. Kim, P. Wang, YH. Li, XL. Liu, S. Octaviani, D. Yang, G. Ma

**Analysis and interpretation of data (e.g., statistical analysis, biostatistics, Computational analysis):** Z. Tian, H. Zhang, A. Pompetti, G. Calendo

**Writing, review, and/or revision of the manuscript:** Z. Tian, H. Zhang, J. Huang

**Administrative, technical, or material support (i.e., reporting or organizing Data, constructing databases):** W. Xu, M. Keough, J. Zhang, H. Zheng, E. Stieglitz, C. Smith, J. Huang

**Study supervision:** J. Huang

## Acknowledgments

We sincerely thank Dr. Peter S. Klein at University of Pennsylvania, Dr. Rona Yaeger at Memorial Sloan Kettering Cancer Center, Dr. Roger K. Strair at Cooper University Health Care and Drs. Jean-Pierre Issa, Xiaoxin (Luke) Chen, Nora Engel, Shumei Song, and Jaroslav Jelinek at Coriell Institute for Medical Research for their insightful comments and discussion. We thank all the members of Huang lab for their help and discussions. We especially thank Drs. Martin Carroll and Alexander Perl at the University of Pennsylvania for his guidance and advice, as well as the University of Pennsylvania Stem Cell & Xenograft Core for providing AML samples and PDX support. AML samples were provided by the Stem Cell and Xenograft Core of the Perelman School of Medicine (RRID: SCR_010035).

## Supplementary Figures

**Figure S1. FLT3 inhibition induces upregulation of *RRM1*, *RRM2*, and *RRM2B* expression in *SPRY3*KO/*FLT3*^ITD^+ AML cells. A.** RNA-seq analysis of *RRM1*, *RRM2*, and *RRM2B* expression in *SPRY3*WT and *SPRY3*KO cells treated with vehicle or gilteritinib. Expression levels are shown as log2 normalized counts. Data are presented as mean ± SD. **B and C.** *SPRY3*KO/*FLT3*^ITD^+ MV4-11 AML cells were treated with AC220 or gilteritinib, followed by quantification of *RRM1*, *RRM2*, and *RRM2B* expression by RT-qPCR; Data represent the means ± SD of three replicates. D and E. *NRAS*^mut^/*FLT3*^ITD+^ MV4-11 AML cells were treated with AC220 or gilteritinib, followed by quantification *of RRM1*, *RRM2*, and *RRM2B* expression by RT-qPCR; Data represent the means ± SD of three replicates.

**Figure S2. Fludarabine, gemcitabine, and hydroxyurea exhibit minimal single-agent cytotoxicity in MV4-11 AML cells. A-C.** Fludarabine, gemcitabine, and hydroxyurea exhibited minimal cytotoxicity as single agents in MV4-11 AML cells. Data represent the means ± SD of three replicates.

**Figure S3.**

**Figure S4. Cell-cycle analysis of *NRAS*^WT^ and *NRAS*^mut^ *FLT3*^ITD^+ AML cells following FLT3 and RNR inhibition. A.** Quantification of the cell-cycle distribution of *NRAS*^WT^, *NRAS*^G12C^, and *NRAS*^Q61K^ AML cells treated with DMSO, gilteritinib, clofarabine, or the combination of gilteritinib and clofarabine. Cells were harvested 24 h after treatment and analyzed by flow cytometry. The percentages of cells in the G0/G1, S, and G2/M phases are shown. Data are presented as mean ± SD from three independent experiments. **B.** Representative flow cytometry histograms showing DNA content and cell-cycle distribution for each treatment group.

**Figure S5. Transcriptomic analysis of *NRAS*-mutant AML cells following gilteritinib and clofarabine treatment. A.** Volcano plots showing differential gene expression between *NRAS*^G12C^ and *NRAS*^WT^ AML cells treated with vehicle, gilteritinib, clofarabine, or the combination of gilteritinib and clofarabine. Red and blue dots indicate significantly upregulated and downregulated genes, respectively. **B.** Heatmap showing differential expression of representative genes involved in RNR-mediated nucleotide biosynthesis, DNA replication, cell-cycle progression, and DNA damage/repair in *NRAS*^G12C^ versus *NRAS*^WT^ AML cells under the indicated treatment conditions. Colors represent log2 fold changes. **C.** Hallmark pathway analysis showing differential pathway activity in *NRAS*^G12C^ versus *NRAS*^WT^ AML cells across the indicated treatment conditions. Dot color represents the pathway enrichment statistic, and gray shading indicates pathways with a Benjamini–Hochberg (BH)-adjusted *P* < 0.05. **D.** Gene set enrichment analysis (GSEA) of the RRM2 126-gene signature, Hallmark E2F Targets, and Hallmark MYC Targets V1 in *NRAS*^G12C^ versus *NRAS*^WT^ AML cells under the indicated treatment conditions. The close correspondence between the *RRM2* signature and the MYC/E2F target gene sets supports a functional link between MYC/E2F signaling and RNR-mediated nucleotide biosynthesis.

**Figure S6. Clofarabine reverses *SPRY3*KO-induced gilteritinib resistance in CDX models of AML. A.** Schematic representation of the MV4-11 CDX *in vivo* study design. **B.** Images of *in vivo* BLI assessment of NSG mice engrafted with *SPRY3*WT and *SPRY3*KO luciferase-tagged MV4-11 cells during the trial. **C and F.** Quantification of BLI data from the (C) *SPRY3*WT and (F) *SPRY3*KO CDX studies. **D and G.** Kaplan-Meier curves representing survival data of subjects from each treatment group in (D) *SPRY3*WT and (G) *SPRY3*KO CDX studies. **E and H.** The combination treatment had no effect on body weight of recipients from both (E) *SPRY3*WT and (H) *SPRY3*KO CDX studies. Data represent the means ± SD (n = 4/group).

**Figure S7. Clofarabine effectively reverses gilteritinib resistance in *NRAS*^Q61K^/*FLT3*^ITD^+ PDX models of AML**. **A.** Measurements of spleen length and weight at the study termination in the gilteritinib-sensitive model. **B.** Representative images of spleen and spleen sections and corresponding immunohistochemical staining for hCD45 from the four treatment groups in the gilteritinib-sensitive model; Scale bars, 100 µm and 50 µm. **C.** Measurements of spleen length and weight at the study termination in the gilteritinib-resistant model.

**Figure S8. Clofarabine restores gilteritinib sensitivity in an *NRAS*-mutant/*FLT3* S451F AML PDX model as measured by hCD45+/hCD33+ leukemia cells.** NSG mice engrafted with a primary AML PDX harboring *NRAS* (Q61L and G12C) and *FLT3* S451F mutations were treated with vehicle, gilteritinib, clofarabine, or the combination of gilteritinib and clofarabine (n = 6 mice per group) for 2 weeks (6 total treatments). At the experimental endpoint, leukemia burden was assessed by flow cytometric analysis of human hCD45+/hCD33+ double-positive leukemia cells in the bone marrow (A, B), spleen (C), and peripheral blood (D). The percentage of hCD45+/hCD33+ cells is shown. Each dot represents an individual mouse, and bars indicate the mean ± SD. Consistent with the results shown in Figure 7, combination treatment significantly reduced leukemia burden compared with vehicle or either monotherapy, with the greatest effect observed in the bone marrow. Statistical significance is indicated as *P* < 0.05, P < 0.01, *P* < 0.001; ns, not significant.

