## Supplementary Figures for "Targeting Ribonucleotide Reductase Overcomes FLT3 Inhibitor Resistance in Acute Myeloid Leukemia"

Figure S1.

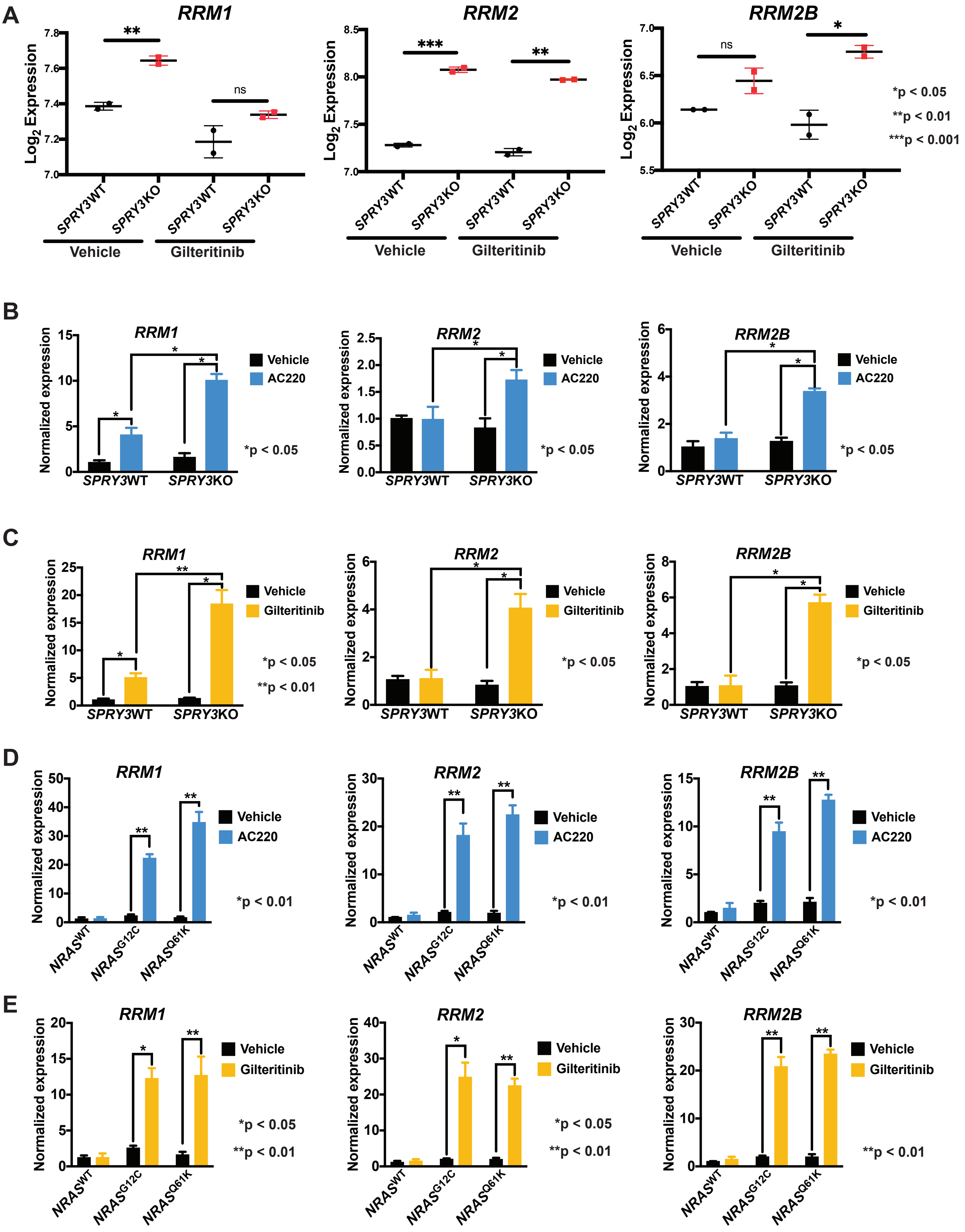


Figure S1

**Figure S2.**

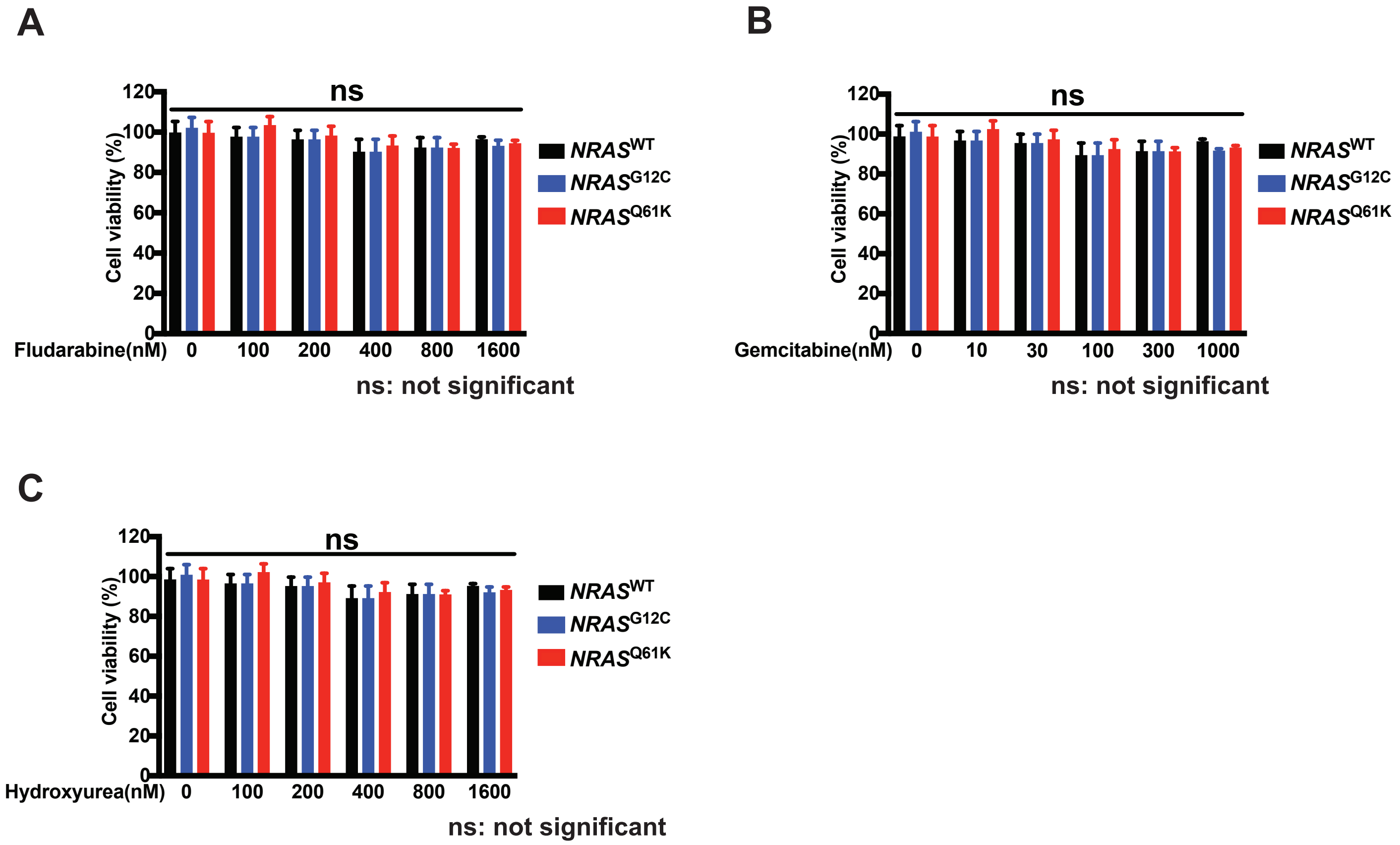
A
ns
Cell viability (%)
120
100
80
60
40
20
0
Fludarabine(nM) 0 100 200 400 800 1600
NRAS^WT
NRAS^G12C
NRAS^Q61K
ns: not significant
B
ns
Cell viability (%)
Gemcitabine(nM) 0 10 30 100 300 1000
NRAS^WT
NRAS^G12C
NRAS^Q61K
ns: not significant
C
ns
Cell viability (%)
Hydroxyurea(nM) 0 100 200 400 800 1600
NRAS^WT
NRAS^G12C
NRAS^Q61K
ns: not significant

Figure S3.

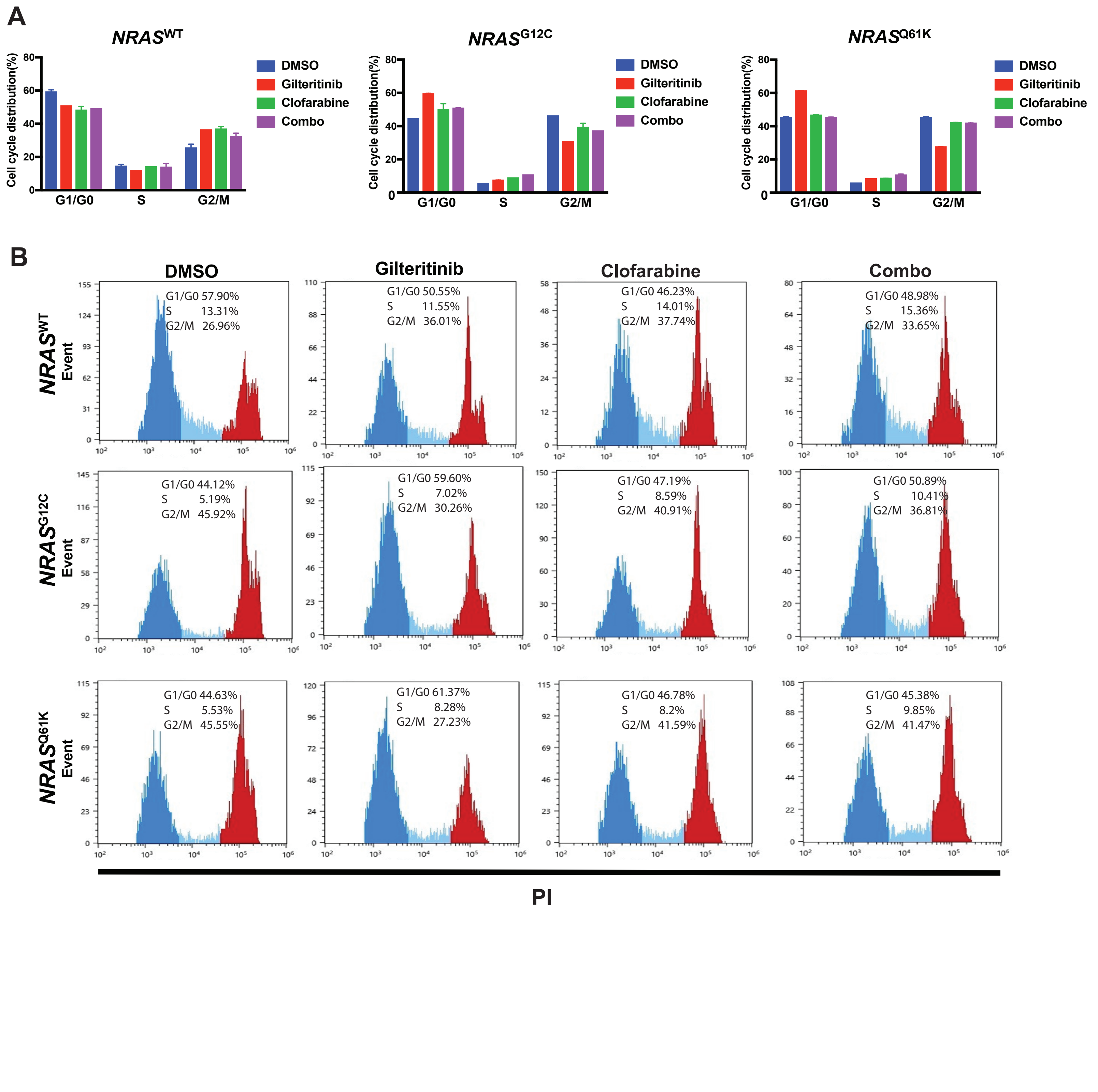
A
NRASWT
NRASG12C
NRASQ61K
Cell cycle distribution(%)
DMSO
Gilteritinib
Clofarabine
Combo
G1/G0
S
G2/M
B
DMSO
Gilteritinib
Clofarabine
Combo
Event
G1/G0 57.90% S 13.31% G2/M 26.96%
G1/G0 50.55% S 11.55% G2/M 36.01%
G1/G0 46.23% S 14.01% G2/M 37.74%
G1/G0 48.98% S 15.36% G2/M 33.65%
G1/G0 44.12% S 5.19% G2/M 45.92%
G1/G0 59.60% S 7.02% G2/M 30.26%
G1/G0 47.19% S 8.59% G2/M 40.91%
G1/G0 50.89% S 10.41% G2/M 36.81%
G1/G0 44.63% S 5.53% G2/M 45.55%
G1/G0 61.37% S 8.28% G2/M 27.23%
G1/G0 46.78% S 8.2% G2/M 41.59%
G1/G0 45.38% S 9.85% G2/M 41.47%
PI

**Figure S4.**

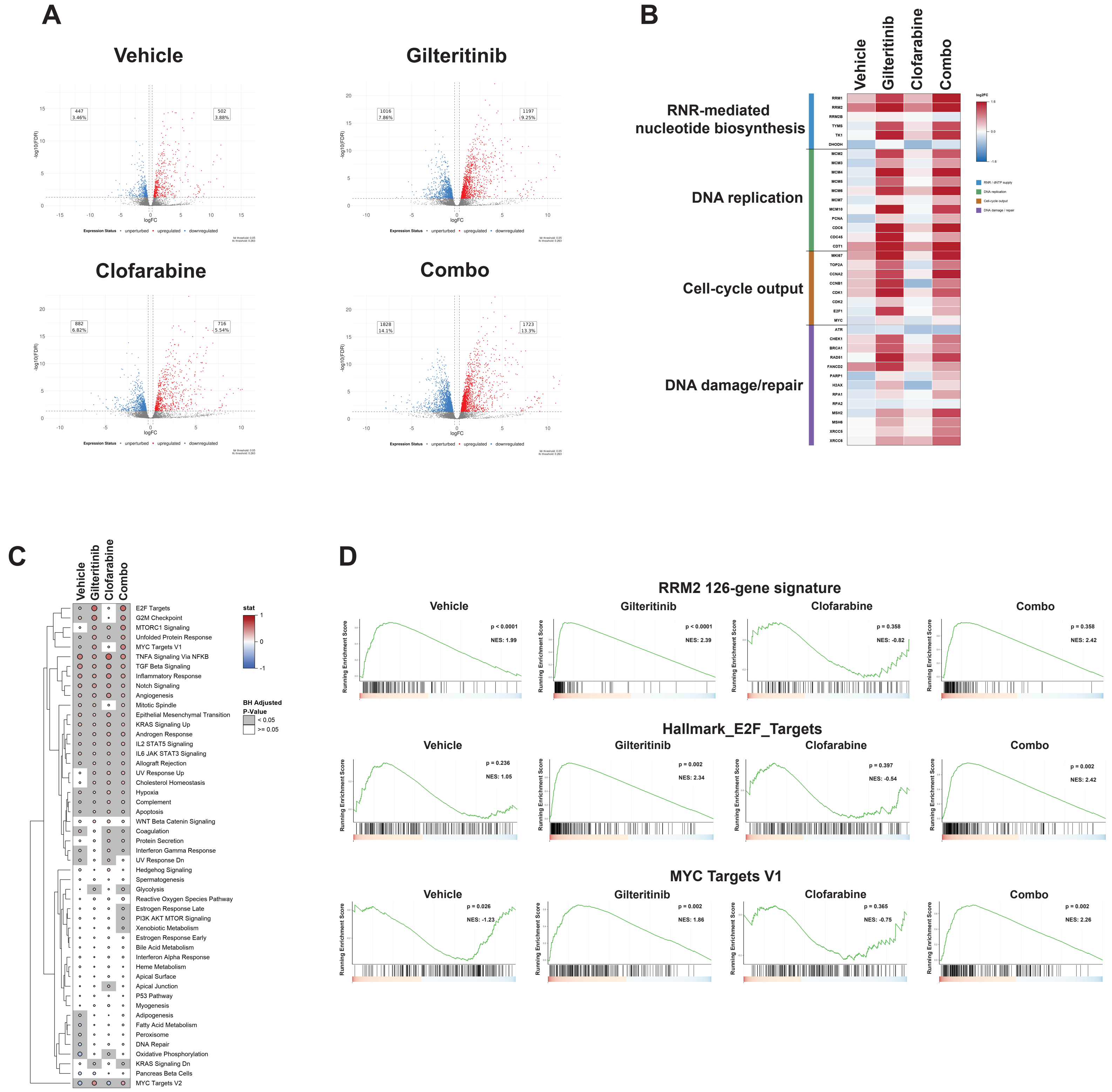


Figure S4

Figure S5.

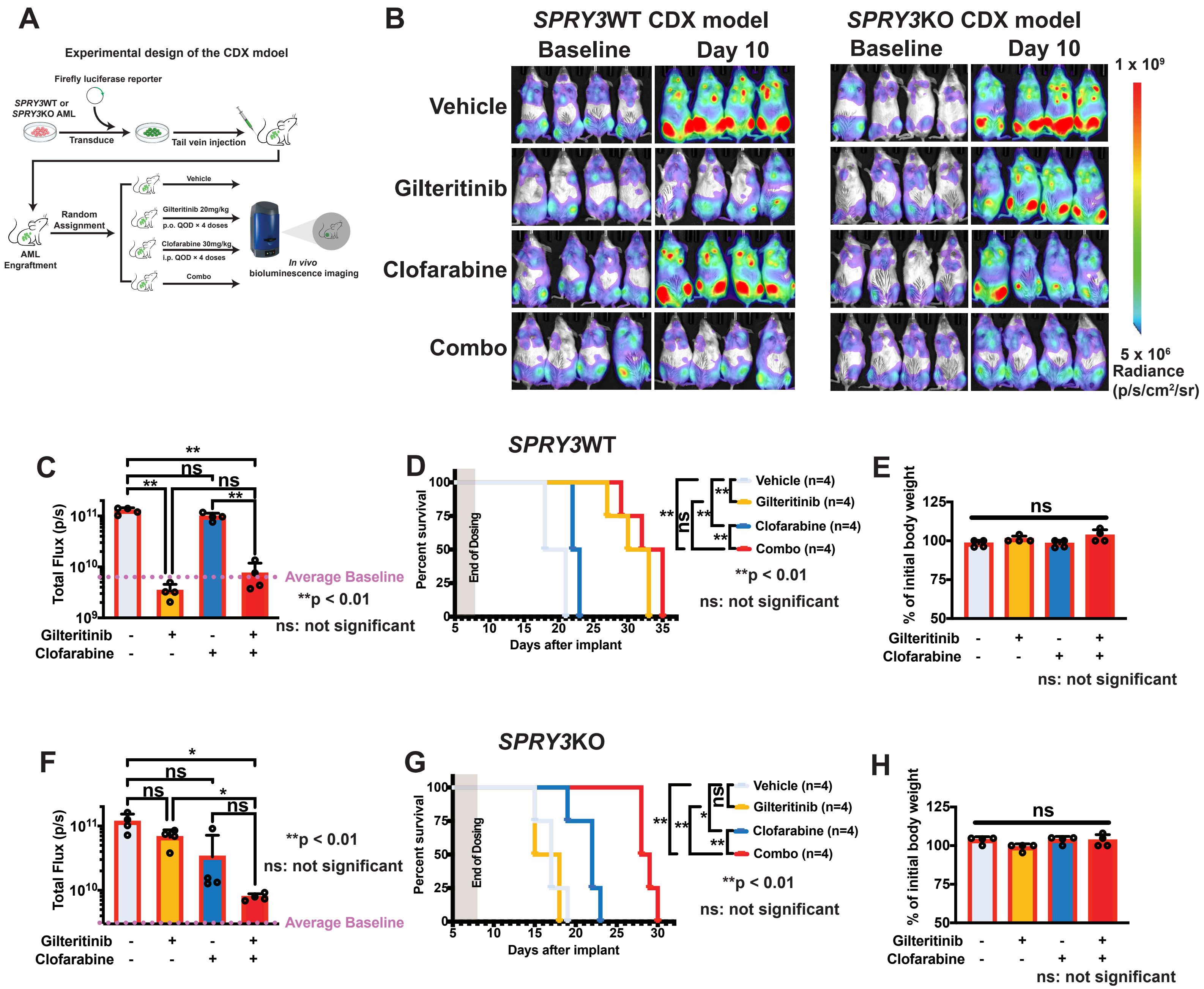

A
Experimental design of the CDX mdoel
Firefly luciferase reporter
SPRY3WT or SPRY3KO AML
Transduce
Tail vein injection
Random Assignment
AML Engraftment
Vehicle
Gilteritinib 20mg/kg p.o. QOD × 4 doses
Clofarabine 30mg/kg i.p. QOD × 4 doses
Combo
In vivo bioluminescence imaging
B
SPRY3WT CDX model
SPRY3KO CDX model
Baseline
Day 10
Vehicle
Gilteritinib
Clofarabine
Combo
1 x 10^9
5 x 10^6
Radiance (p/s/cm²/sr)
C
Total Flux (p/s)
Average Baseline
**p < 0.01
ns: not significant
Gilteritinib
Clofarabine
D
SPRY3WT
Percent survival
End of Dosing
Days after implant
Vehicle (n=4)
Gilteritinib (n=4)
Clofarabine (n=4)
Combo (n=4)
**p < 0.01
ns: not significant
E
% of initial body weight
ns
ns: not significant
F
Total Flux (p/s)
Average Baseline
**p < 0.01
ns: not significant
G
SPRY3KO
Percent survival
End of Dosing
Days after implant
Vehicle (n=4)
Gilteritinib (n=4)
Clofarabine (n=4)
Combo (n=4)
**p < 0.01
ns: not significant
H
% of initial body weight
ns
ns: not significant

**Figure S6.**

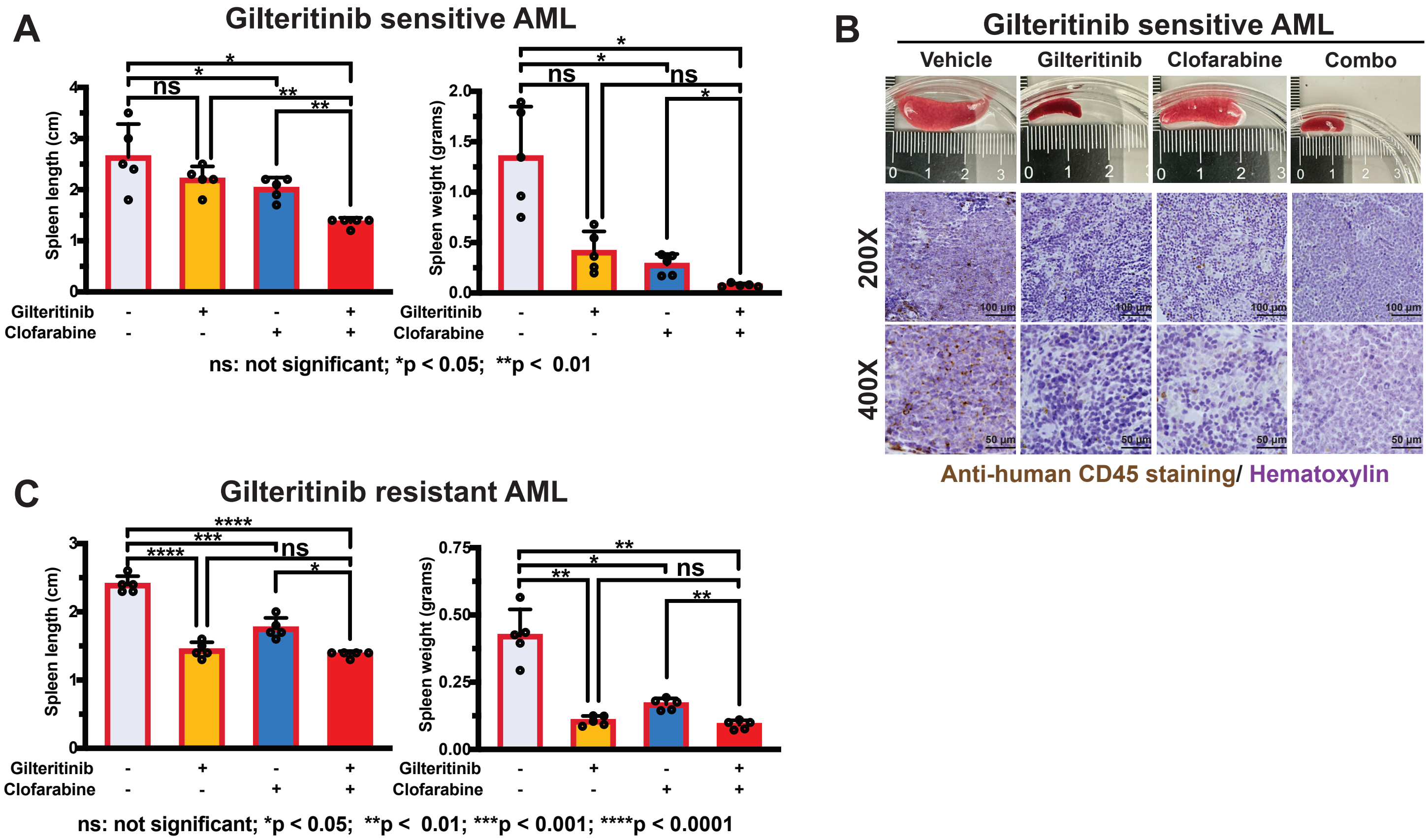
A
Gilteritinib sensitive AML
Spleen length (cm)
Spleen weight (grams)
ns
Gilteritinib
Clofarabine
ns: not significant; *p < 0.05; **p < 0.01
B
Gilteritinib sensitive AML
Vehicle
Gilteritinib
Clofarabine
Combo
200X
400X
Anti-human CD45 staining/ Hematoxylin
C
Gilteritinib resistant AML
ns: not significant; *p < 0.05; **p < 0.01; ***p < 0.001; ****p < 0.0001

**Figure S7.**

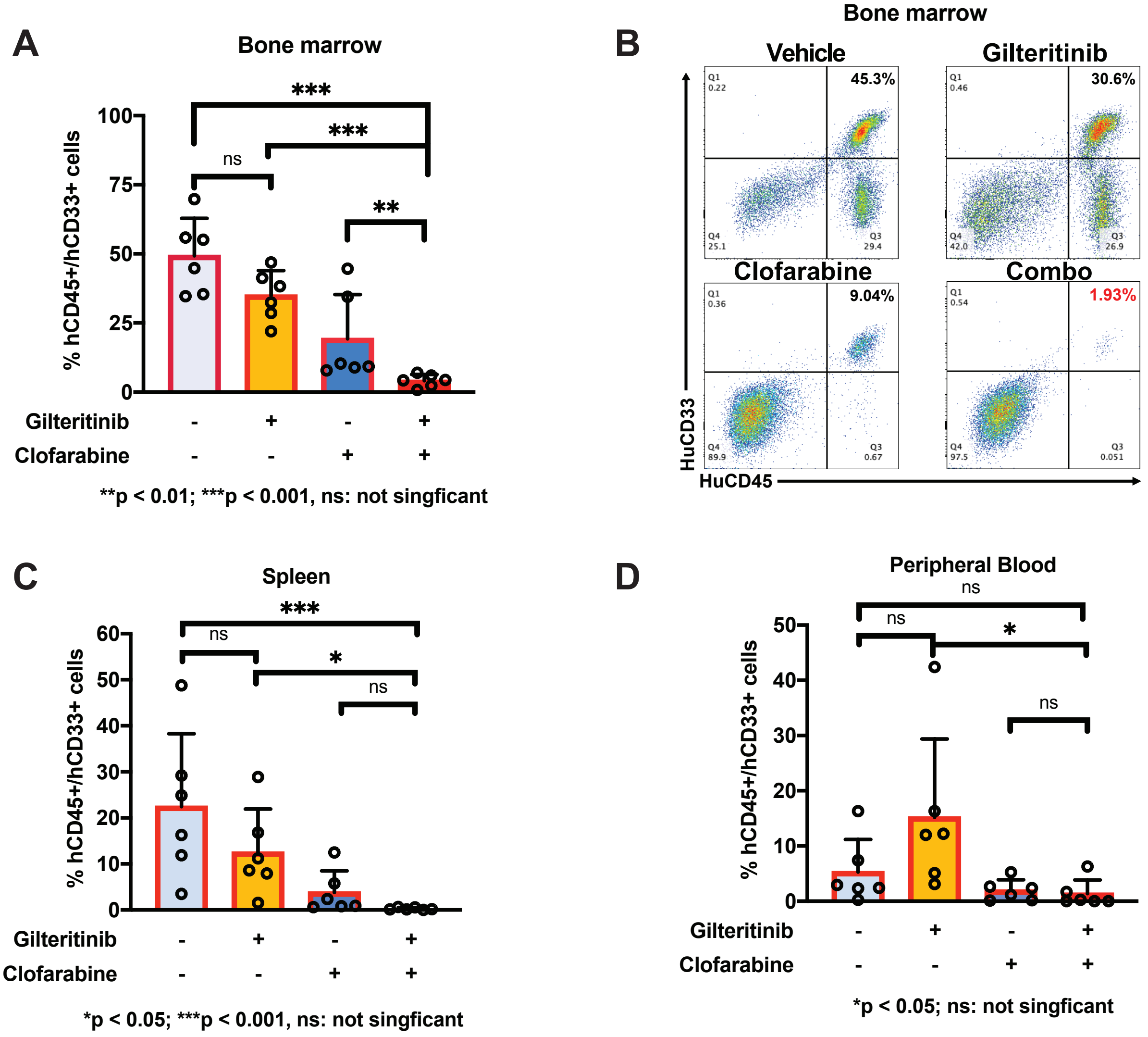
A
Bone marrow
% hCD45+/hCD33+ cells
100
75
50
25
0
ns
***
***
**
Gilteritinib - + - +
Clofarabine - - + +
**p < 0.01; ***p < 0.001, ns: not singficant
B
Bone marrow
Vehicle
Gilteritinib
45.3%
30.6%
Clofarabine
Combo
9.04%
1.93%
HuCD33
HuCD45
C
Spleen
% hCD45+/hCD33+ cells
60
50
40
30
20
10
0
***
ns
*
ns
Gilteritinib - + - +
Clofarabine - - + +
*p < 0.05; ***p < 0.001, ns: not singficant
D
Peripheral Blood
ns
ns
*
ns
% hCD45+/hCD33+ cells
50
40
30
20
10
0
Gilteritinib - + - +
Clofarabine - - + +
*p < 0.05; ns: not singficant
